# Identification of genetic loci in lettuce mediating quantitative resistance to fungal pathogens

**DOI:** 10.1101/2022.03.08.483472

**Authors:** Harry Pink, Adam Talbot, Abi Graceson, Juliane Graham, Gill Higgins, Andrew Taylor, Alison C. Jackson, Maria Truco, Richard Michelmore, Chenyi Yao, Frances Gawthrop, David Pink, Paul Hand, John P. Clarkson, Katherine Denby

**Affiliations:** Centre for Novel Agricultural Products (CNAP), Biology Department, University of York, Wentworth Way, York, YO10 5DD; Department of Agriculture and Environment, Harper Adams University, Newport, Shropshire, TF10 8NB; School of Life Sciences, University of Warwick, Wellesbourne Campus, Warwick, CV35 9EF; Genome Center, University of California Davis, One Shields Ave, Davis, California, USA, 95616; A. L. Tozer Ltd., Pyports, Downside Road, Cobham, Surrey, KT11 3EH

**Author notes:** Corresponding author: Katherine Denby. These authors contributed equally to this research (Harry Pink, Adam Talbot).

**Keywords:** lettuce, disease resistance, quantitative genetics, *Botrytis cinerea*, *Sclerotinia sclerotiorum*, transcriptomics

## Abstract

*Lactuca sativa* L. (lettuce) is an important leafy vegetable crop grown and consumed globally. Chemicals are routinely used to control major pathogens, including the causal agents of grey mould (*Botrytis cinerea*) and lettuce drop (*Sclerotinia sclerotiorum*). With increasing prevalence of pathogen resistance to fungicides and environmental concerns, there is an urgent need to identify sources of genetic resistance to *B. cinerea* and *S. sclerotiorum* in lettuce. We demonstrated genetic variation for quantitative resistance to *B. cinerea* and *S. sclerotiorum* in a set of 97 diverse lettuce and wild relative accessions, and between the parents of lettuce mapping populations. Transcriptome profiling across multiple lettuce accessions enabled us to identify genes with expression correlated with resistance, predicting the importance of post-transcriptional gene regulation in the lettuce defence response. We identified five genetic loci influencing quantitative resistance in a F10 mapping population derived from a *Lactuca serriola* (wild relative) x lettuce cross, which each explained 5–10% of the variation. Differential gene expression analysis between the parent lines, and integration of data on correlation of gene expression and resistance in the diversity set, highlighted potential causal genes underlying the quantitative trait loci.

**Key Message:** We demonstrate genetic variation for quantitative resistance against important fungal pathogens in lettuce and its wild relatives, map loci conferring resistance and predict key molecular mechanisms using transcriptome profiling.

## Introduction

*Lactuca sativa* L. (lettuce) is an economically valuable leafy vegetable with production worth more than £200 million in the UK (Defra Horticulture Statistics, 2020) and $2.4 billion in the USA (USDA-NASS, 2019). Lettuce is susceptible to a wide range of plant pathogens including the fungal necrotrophs *Botrytis cinerea* Pers. and *Sclerotinia sclerotiorum* (Lib.) de Bary, the causal agents of grey mould and lettuce drop, respectively. *B. cinerea* was ranked second for fungal pathogens of scientific and economic importance (Dean et al. 2012) while up to 50% of lettuce yields may be lost due to *S. sclerotiorum* (Young et al. 2004). Chemical control is routinely used but there is an urgent need to identify sources of host genetic resistance given the costs of preventative pesticide sprays, the prevalence of fungicide-resistant isolates of both pathogens in the field (Zhou et al. 2014; Rupp et al. 2017; Hou et al. 2018) and the increasing withdrawal of approved fungicides though legislation.

Pathogens with a biotrophic lifestyle parasitize and extract nutrients from living plant tissue, whereas necrotrophic pathogens rapidly kill their host, extracting nutrients from the dead tissue. A plant’s response to infection varies depending on the pathogen lifestyle (Mengiste 2012). Complete disease resistance against specific isolates of biotrophic pathogens is often seen, conferred by a single dominant host gene. Many of these genes encode nucleotide binding leucine rich-repeat (NLR) proteins, which directly or indirectly detect the presence of pathogen effectors (virulence factors delivered into host cells to aid infection)(Cui et al. 2013). Similarly, resistance to host-specific necrotrophic pathogens, such as *Cochliobolus carbonum*, the causal agent of northern leaf spot in maize, is controlled by single gene traits conferring toxin sensitivity on susceptible host plants (Panaccione et al. 1992). In contrast, resistance to broad-host range necrotrophic pathogens (of which *B. cinerea* and *S. sclerotiorum* are prime examples) is usually a quantitative trait, with a continuum of phenotypes rather than two distinct classes of resistant and susceptible (Corwin and Kliebenstein 2017). This quantitative resistance is controlled by multiple genes with small to moderate effects (Roux et al. 2014).

Molecular analyses, mostly in the model plant Arabidopsis, have identified numerous components of the plant response to infection by *B. cinerea* (AbuQamar et al. 2017) and *S. sclerotiorum* (Wang et al. 2019) incorporating pathogen detection, signal transduction and activation of host defences. For many of these individual components, mutants (knock-outs or overexpressors) have been used to assess their impact on disease outcome. However, these single-gene transgenic studies are much more difficult in non-model plants and do not help us understand the genetic variation in natural or managed populations, nor the relative contribution different genes/loci make to overall plant resistance. Genetic loci contributing to quantitative disease resistance (QDR) against *B. cinerea* have been mapped in a number of different plant species including Arabidopsis (Denby et al. 2004; Rowe and Kliebenstein 2008; Coolen et al. 2019), tomato (Finkers et al. 2007; Szymański et al. 2020), *Brassica rapa* (Zhang et al. 2016) and *Gerbera hybrida* (Fu et al. 2017). Multiple studies investigating QDR against *S. sclerotiorum* in *B. napus*, sunflower and soybean using quantitative trait loci (QTL) mapping and genome-wide association studies (GWAS) (reviewed in (Wang et al. 2019) have identified many loci, each with a minor effect on QDR. However, what is lacking is knowledge of the molecular mechanisms underlying these loci. Recombination frequency within mapping populations and linkage disequilibrium in association panels typically limit resolution of the loci. Co-localisation of genetic loci for different traits (such as those mediating the accumulation of specific metabolites with QTL controlling QDR against *B. cinerea*) can be informative in predicting causal genes or mechanisms. For example, camalexin accumulation QTL co-localised with QTL influencing lesion size after *B. cinerea* infection in Arabidopsis (Rowe and Kliebenstein 2008) and QTL in *B. rapa* controlling the accumulation of glucosinolates were co-localised with *B. cinerea* resistance QTL (Zhang et al. 2016). Szymánski et al. (Szymanski et al. 2020) combined metabolic QTL and expression QTL with QTL mediating tomato fruit resistance to *B. cinerea* to predict specific flavonoids important for host resistance.

In lettuce, QTL mapping has been used extensively to characterise dominant resistance phenotypes against the oomycete pathogen *Bremia lactucae*, which causes downy mildew. More than 30 downy mildew resistance genes have been identified (Parra et al. 2016, 2021). However, mapping of genetic determinants of QDR in lettuce against *B. cinerea* or *S. sclerotiorum* is in its infancy. Recently, two QTL were reported for field resistance to lettuce drop in the Reine des Glaces × Eruption mapping population (Mamo et al. 2019). However, lettuce drop can be caused by both *S. sclerotiorum* and *Sclerotinia minor* and in this case the fields were inoculated with *S. minor*, which has a different infection strategy (infection via mycelia in the soil) than *S. sclerotiorum* (infection via germinating ascospores). To our knowledge no lettuce QTL have been reported for QDR against *B. cinerea* or *S. sclerotiorum*.

Here we demonstrate genetic variation in resistance to *B. cinerea* and *S. sclerotiorum* in a lettuce diversity set (Walley et al. 2017) including *L. sativa* cultivars and wild relatives and exploit a bi-parental mapping population to identify QTL mediating resistance to both pathogens. Transcriptome profiling of a selection of the diversity set lines identified genes with expression correlated with disease resistance and highlighted post-transcriptional gene regulation (in particular, gene silencing) and pathogen recognition as determinants of resistance. Moreover, we integrated the diversity set and mapping population parent line transcriptome data to predict causal genes underlying the QTL.

## Methods

### Lettuce lines and plant growth

The Diversity Fixed Foundation Set (DFFS) comprises 96 lettuce accessions selected from the lettuce collection at the UK Vegetable Genebank, Wellesbourne, UK and the international *Lactuca* collection at the Centre for Genetic Resources, Netherlands. The set includes 17 wild species as well as a range of cultivated varieties (Walley et al. 2017). For detached leaf inoculation assays, two lettuce seeds for each line were sown into 9 cm x 9 cm x 10 cm plastic pots filled with well-packed Levington’s M2 growing media (Harper Adams University) or into 56 mm x 56 mm x 50 mm plug plant cells filled with well-packed Levington’s F2S growing media (Universities of Warwick and York). The seeds were covered with a thin layer of vermiculite and watered frequently to ensure the growing media remained damp. Following germination, seedlings were thinned to one per pot. Plants were grown in a glasshouse with supplemental lighting provided for 16 hours and heating set to 18°C. Experiments were carried out at Harper Adams University (52°46’46.02”N, 2°25’37.68”W), University of Warwick (52°12’37.31”N, 1°36’0.42”W) and University of York (53°56’44.16”N, -1°03’28.44”W) between 2015 and 2017 (Table 2).

### Infection assays

The *S. sclerotiorum* isolate L6 was used in all experiments (Taylor et al. 2018). Sclerotia and apothecia were produced and ascospores captured onto filter paper as described by (Clarkson et al. 2014). Spore suspensions were prepared by agitating a section of filter paper in 5 mL distilled water until the water appeared cloudy. The suspension was filtered through two layers of miracloth and diluted to 5×10^5^ spores per mL following counts using a haemocytometer. *B. cinerea* inoculum (pepper isolate)(Denby et al. 2004) was prepared by inoculating sterile tinned apricot halves. The inoculated apricot halves were sealed in Petri dishes and left in the dark at 25°C for 14 days to facilitate sporulation. Spore suspensions of *B. cinerea* were prepared by washing off conidiospores in 3 mL distilled water and filtering through two layers of miracloth. The suspension was again diluted to 5×10^5^ spores per mL.

The third leaf from four-week-old lettuce plants was removed and placed on 0.8% (w/v) agar in sealable propagator trays. Leaves that were damaged were discarded. Each leaf was inoculated with a 5 µL drop of either *S. sclerotiorum* or *B. cinerea* spore suspension on either side of the mid-vein. The trays of leaves were covered to maintain humidity and placed in a controlled environment cabinet at 22°C, 80% humidity and a cycle of 12 h light:12 h dark. Overhead photographs were taken of each tray between 48 h and 72 h post-inoculation (hpi) including a scale bar to enable measurement of lesion area using ImageJ2. Overlapping lesions, lesions that had spread to the leaf edge or lesions that had failed to initiate were not measured. For assessment of the complete diversity set one leaf (two inoculation sites) of each lettuce line was included in a single experiment, and experiments were repeated for ten consecutive weeks. To compensate for differences between experimental replicates and missing lesions, a REML was used to identify sources of variation between square root lesion size. Means were predicted using the least-squares method.

For the assessment of lesion size coupled to gene expression, sixteen lines were used in experiments with both pathogens, with a further 10 lines inoculated with a single pathogen (five with *B. cinerea*; five with *S. sclerotiorum*). Detached leaf infection assays were carried out using three replicate leaves of each accession for each pathogen (apart from eight varieties with two replicates for *S. sclerotiorum* infection and four varieties with two replicates for *B. cinerea* infection) giving 114 leaves in total. Lesion size was measured at 42 hpi for *S. sclerotiorum* and 46 hpi for *B. cinerea* (Supplementary Dataset 3) with leaves harvested at 43 hpi (*S. sclerotiorum)* and 48 hpi (*B. cinerea)* for transcriptome profiling.

For assessment of lesion size in the mapping population in each replicate 120 lines were grown to four weeks old before leaf 3 was detached and placed on agar prior to droplet inoculation. This was repeated in a randomised design to a total of eight experiments. Mean lesion size per line was estimated using a least squares method.

### Polytunnel Assay

Whole-head lettuce disease assays for both *B. cinerea* and *S. sclerotiorum* were performed in duplicate at the University of Warwick and Harper Adams University. Eighteen plants per accession (per pathogen inoculation) were grown to 4 weeks old in the glasshouse (as above) before being transplanted to 24 cm diameter pots in Levington’s F2S soil. Plants were placed in a blocked randomisation design within a sealed polytunnel. Spores were collected as above in 500 mL sterile distilled water and diluted to 1×10^5^ spores/mL before being sprayed directly onto each plant using a hand-sprayer. Plants were sprayed with inoculum until saturation and inoculum run off, and irrigated via spray irrigation from 1.5 m tall irrigators every 2 hours from 6 am until 2 am the next day, for 10 minutes at each interval. Plants were assessed twice weekly, with the following scale: 0 – no symptoms, 1 – visible lesions on lower leaves, 2 – visible lesions on majority of leaves, 3 – severe disease symptoms over entire plant, 4 – total plant collapse. Plants were monitored to 6 weeks post-transplanting for disease symptoms. Area under the disease progression curve (AUDPC) was calculated using the trapezoid rule with the *Agricolae* R package. ANOVA was used to determine significant differences in variation followed by a Tukey HSD test to determine significant differences between varieties.

### QTL analysis

The RIL population of 234 F6 lines generated from crossing the Armenian *L. serriola* 999 and *L. sativa* PI251246, including genotyping of the population and generation of a genetic map using 2677 markers, was previously described (Han et al. 2021). Minimal linkage disequilibrium was observed, and QTL analysis was performed in the R/qtl package (Broman et al. 2003). Recombination fraction was estimated using the est.rf function. The least-squares predicted mean of square-root lesion size (mm^2^) was used as the phenotyping score for each RIL. ‘calc.genoprob’ was used (with an error probability of 0.001 and a step-limit of 2 cM), which utilises hidden Markov models to estimate true underlying genotype between markers. A single QTL scan was performed, using ‘scanone’ as a preliminary measure using Haley-Knott regression (Haley and Knott 1992), for which the significance threshold was calculated by a permutation test of 1,000 imputations with an alpha value of 0.05. A search for epistatic interactions between loci was conducted using ‘scantwo’ with significance calculated based on a permutation test (750 imputations). QTL above the permutation threshold for either algorithm were then fitted to the multi-QTL model selection pipeline using ‘makeqtl’ and ‘fitqtl’. Percentage variance explained statistics were calculated by ‘fitqtl’. Once fitted, ‘addqtl’ was used to search for additional QTL missed by the preliminary scan. Peak QTL positioning was further adjusted using ‘refineqtl’. Finally, a forward/backwards selection model was applied with ‘stepwiseqtl’ to give the final QTL, with model penalties calculated at an alpha level of 0.05 (Manichaikul et al. 2009). Haley-Knott regression (Haley and Knott 1992) was used for the model selection stages. Epistatic interactions between the QTL loci were passed to the ‘stepwiseqtl’ algorithm, but none passed the significance threshold. To identify confidence regions surrounding each QTL, ‘lodint’ was used, a size of 1.5 Logarithm of the odds score (LOD) around each QTL peak was selected and expanded to the next marker. Flanking markers of the 1.5LOD confidence interval were mapped back to the *Lactuca sativa* cv. Salinas v8 genome (Reyes-Chin-Wo et al. 2017) to identify genes located within the QTL. Another R package, LinkageMapView was used to visualise the genetic map (Ouellette et al. 2017).

### Gene Expression profiling

Leaves were infected with *S. sclerotiorum* (Taylor et al. 2017) or *B. cinerea* as per the detached leaf assays outlined above, and samples were harvested using a size 6 cork borer centred on the lesion. All expression profiling experiments used *B. cinerea* pepper isolate (Windram et al. 2012). For *S. sclerotiorum* the L6 isolate was used for the diversity set and mapping population parent expression analysis, with the P7 isolate used for response to infection expression (mock vs. inoculated). Infected tissue was snap frozen in liquid nitrogen before RNA extraction using Trizol (Thermo Fisher Scientific) with a lithium chloride purification. Sequencing libraries were prepared using the Illumina TruSeq RNA V2 kit and sequenced using a HiSeq 2500 generating 100 bp paired-end reads or HiSeq 4000 with 75 bp paired-end reads. Read quality was checked with FastQC (Wingett and Andrews 2018). Sequencing reads were aligned to a combined lettuce**–**pathogen transcriptome using Kallisto (Bray et al. 2016). Counts were summarised at the gene level and differential expression analysis was performed using the Limma-Voom pipeline (Law et al. 2014) with a threshold of ≥1.2 log2 fold change. Principal component analysis was performed on gene counts using the “prcomp” function in R and Euclidean distance between samples was used for hierarchical clustering.

### Lesion Size-Gene Expression Correlation

Diversity set reads aligned to lettuce genes were normalised using the trimmed mean of M-values from edgeR (Robinson et al. 2010). Genes with low expression (under 50 counts) and transcripts with low variance (<1.2 logFC between samples) were removed. Spearman correlations were calculated for the relationship between trimmed mean of M expression values and square-root lesion size for each gene (23,111 in *S. sclerotiorum* and 23,164 in *B. cinerea)*. Correlation p-values were calculated using *cor*.*test* in R and corrected using the Benjamini-Hochberg method (Benjamini and Hochberg 1995).

### Gene-Ontology (GO) Enrichment Analysis

GO enrichment analysis was performed for genes whose expression significantly correlated with increased pathogen resistance or susceptibility using the *org*.*At*.*tair*.*db* and *clusterprofiler* R packages (Wu et al. 2021). Arabidopsis orthologs are taken from (Reyes-Chin-Wo et al. 2017).

## Results

### Lettuce accessions vary in lesion development after inoculation with S. sclerotiorum or B. cinerea

A detached leaf assay was previously developed for infection of Arabidopsis by *B. cinerea* and shown to be effective in quantitative assessment of lesion development and correlated with fungal growth within the leaf (Windram et al. 2012). We had also used a similar assay to evaluate susceptibility to *S. sclerotiorum* in multiple Brassica lines (Taylor et al. 2017). We adapted these assays to measure lesion development in lettuce after inoculation with the two closely related necrotrophic fungal pathogens (*B. cinerea* and *S. sclerotiorum*) to assess susceptibility to disease. Using these quantitative assays, we measured lesion size in 97 lettuce accessions (96 lines comprising a Diversity Fixed Foundation set (Walley et al. 2017) plus the cultivar Lolla Rossa) after inoculation with *B. cinerea* or *S. sclerotiorum* spores (Supplementary Dataset 1). Crucially, although these assays use detached leaves, they use spores (ascospores of *S. sclerotiorum* and conidiospores of *B. cinerea*) to mimic the type of infection occurring naturally. This contrasts with the commonly used *S. sclerotiorum* inoculation method of mycelial plugs (Joshi et al. 2016; Barbacci et al. 2020; Chittem et al. 2020). With occasional lack of plant growth and/or lack of infection from the droplet, the average number of lesions (and leaves) per lettuce line inoculated with *B. cinerea* was 16 (eight leaves) and for *S. sclerotiorum* 17 (seven leaves). Restricted Maximum Likelihood (REML) least-squares predicted mean lesion size was calculated for each lettuce accession to account for variation between replicate experiments.

The lettuce diversity set clearly contains genetic variation for susceptibility to *B. cinerea* and *S. sclerotiorum*, as judged by lesion development on detached leaves (Figure 1). Lettuce lines exhibited significant variation in lesion size (p <0.001 for both pathogens), which likely reflects the ability of the pathogens to grow, and hence the effectiveness of the plant defence response to combat infection. Lesion size varied by lettuce type; for example, Iceberg lettuces were significantly more resistant to *B. cinerea* and *S. sclerotiorum* than Butterhead and Cutting types (Supplementary Figure 1). Different lettuce types have large architectural differences (Walley et al. 2017) which alters the ability of pathogens to infect the plant in the field; for example, a more open structure reduces moist environments for spore collection and germination. A benefit of the detached leaf assay is that it identifies architecture-independent sources of resistance that could be deployed across multiple lettuce types. Wild relatives of lettuce (*L. virosa, L. saligna* and *L. serriola)* were significantly more resistant to both *S. sclerotiorum* and *B. cinerea* than the cultivated *L. sativa* (Tukey HSD p<0.05) (Figure 2), suggesting that alleles conferring quantitative resistance against these fungal pathogens may have been lost during the domestication of lettuce.

**Figure 1:**
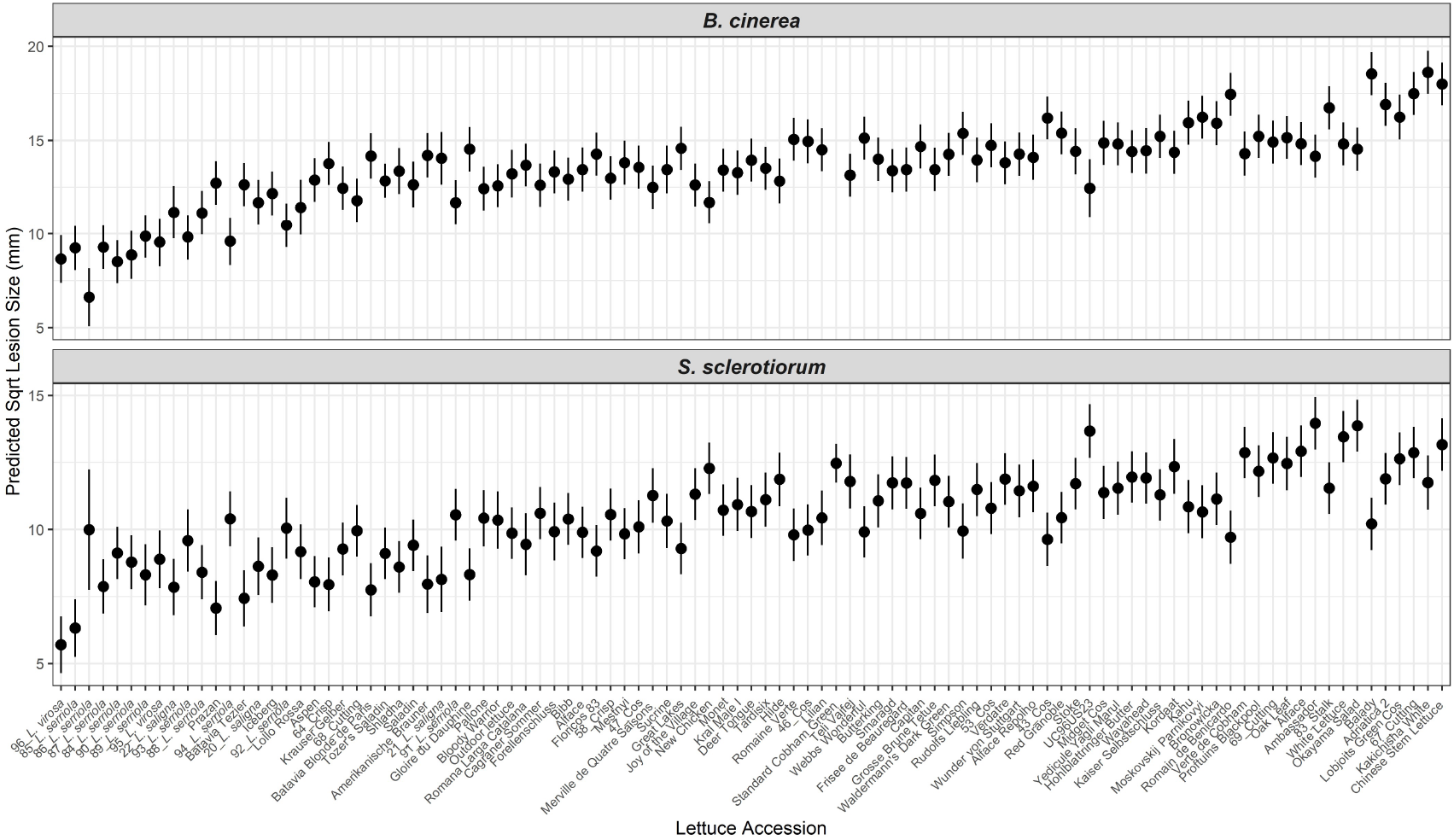
Variation in lesion size after inoculation with *Botrytis cinerea* or *Sclerotinia sclerotiorum* in a set of lettuce accessions. Least-squares REML predicted mean square root of lesion size 64 hours after *Botrytis cinerea* (top) or *Sclerotinia sclerotiorum* (bottom) inoculation of detached lettuce leaves is shown in ascending order of mean lesion size across the two pathogens. Error bars are REML standard error. Lesions measured per accession per pathogen ranges from two (#86 *L. serriola* – *S. sclerotiorum*) and 213 (Tozer Saladin – *B. cinerea*) with a median n=16.

**Figure 2:**
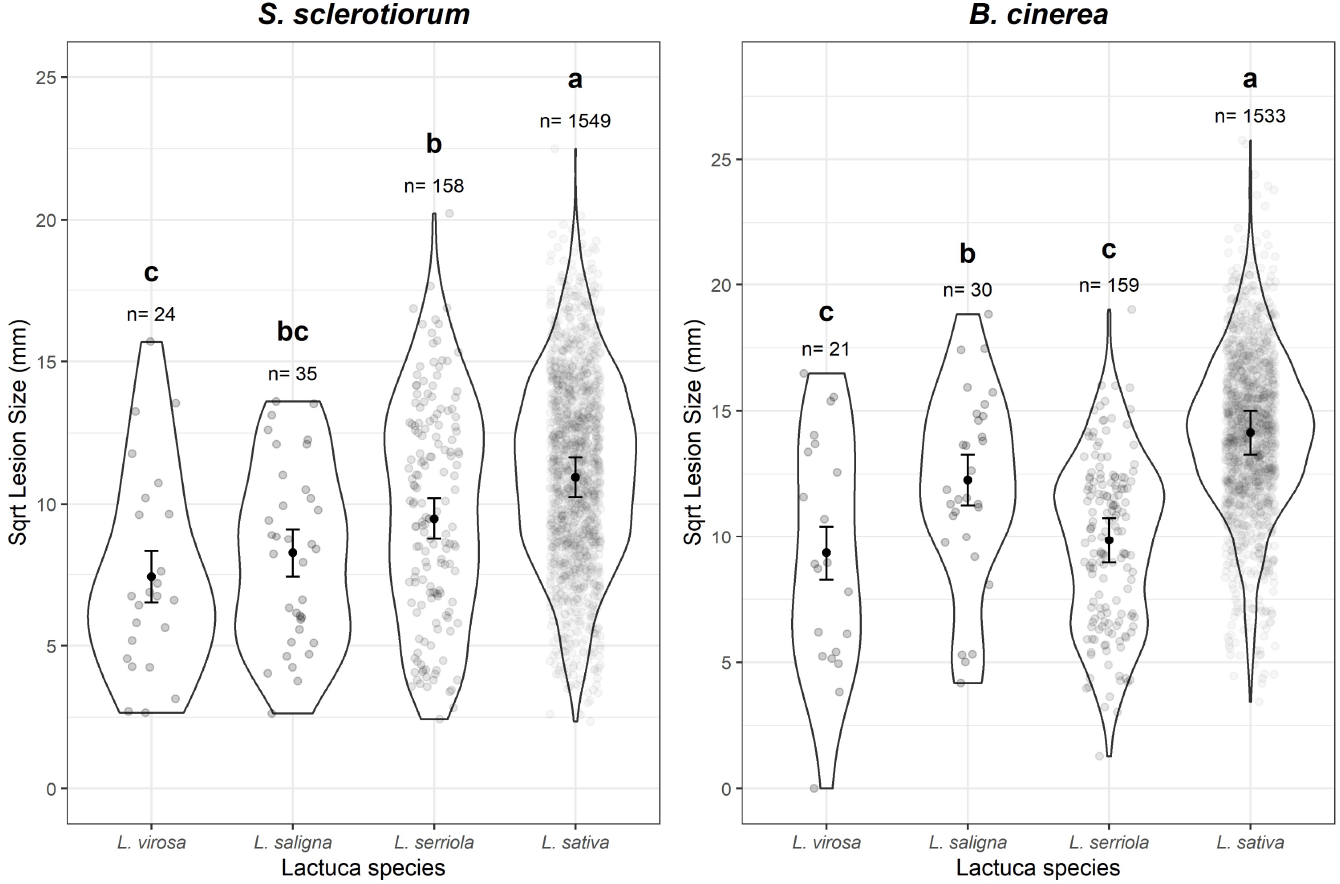
Variation in lesion size after inoculation with *Botrytis cinerea* or *Sclerotinia sclerotiorum* in different lettuce species. Square root lesion size 64 hours after *Sclerotinia sclerotiorum* (left) or *Botrytis cinerea* (right) inoculation of detached lettuce leaves is shown. Grey circles represent individual measured lesions, with areas within the lines showing the distribution of data points. Black circles are the REML predicted mean lesion size per species (correcting for random variation between experimental replicates) with error bars showing REML predicted standard error. Letters represent Tukey post-hoc significance groupings (p< 0.05) performed on the REML model. n is the number of lesions measured from each species.

*B. cinerea* and *S. sclerotiorum* are closely related necrotrophic fungal pathogens sharing many genes and virulence strategies (Amselem et al. 2011), although *B. cinerea* contains a higher number and diversity of genes involved in secondary metabolism (e.g. the production of plant toxins). Consistent with their similarity, we found a correlation (R=0.47, p= 1.1E-6) across the diversity set between lesion size after inoculation with each of the two pathogens (Supplementary Figure 2), raising the prospect of identifying lettuce alleles conferring quantitative resistance against both pathogens.

We performed a whole-head lettuce inoculation experiment to determine whether such an assay could be used in a quantitative manner and if the detached leaf assay data were relevant to whole plants. Four-week-old plants of seven lettuce accessions were sprayed with spore suspensions of *S. sclerotiorum* or *B. cinerea* and humidity was kept high through regular mist irrigation. A disease score was captured for each plant from 14 to 49 days post inoculation, and the AUDPC was calculated to quantify disease symptoms over time (Supplementary Dataset 2). Plots of disease score progression for each accession are shown in Supplementary Figure 3. The relationship between AUDPC from the whole-head assay and lesion size in the detached leaf assay for the seven selected lettuce accessions is shown in Supplementary Figure 4. There is a positive trend between the two measurements: accessions with a higher whole plant disease score (AUDPC) also had a higher detached leaf assay lesion size in response to *B. cinerea* (RS=0.64) and *S. sclerotiorum* (RS=0.61). However, neither correlation showed statistical significance. Notably, the accession with the smallest lesion size in the detached leaf assay (*L. virosa*, line 96) and the accessions with the largest lesion size (Okayama Salad, Ambassador) showed a clear difference in the whole plant assay suggesting that the detached leaf assay phenotypes do have relevance to whole head disease progression. However, the whole plant assay appears unable to distinguish varieties with intermediate levels of resistance. Hence, we proceeded with the detached leaf assay as it provides quantitative measurements over a wider range of values and is carried out under more controlled conditions.

### Diversity set transcriptome profiling identifies gene expression correlated to disease progression

We selected accessions from the lettuce diversity set that exhibited a wide range of lesion size after infection with both pathogens whilst focusing on *L. sativa* varieties to ease transcriptome analysis. The Armenian *L. serriola* line was also included as it is a parent of a key mapping population. Despite a lower number of replicates than in the full diversity set experiment, capturing the lesion size from the exact leaves used for transcriptome analysis enables us to directly link gene expression and lesion size in the same leaf.

A disc of tissue around each developing lesion was sampled for RNAseq transcriptome profiling with three biological replicates for each accession/pathogen combination. Information on the number and mapping of raw reads are in Supplementary Dataset 3. Reads were mapped to a combined *S. sclerotiorum-* or *B. cinerea-*lettuce (*L. sativa* var. Salinas) transcriptome (Derbyshire et al. 2017; Kan et al. 2017; Reyes-Chin-Wo et al. 2017). Approximately 25,000 lettuce genes were present in at least one sample (Supplementary Dataset 4). As lesion size reflects pathogen growth *in planta*, the percentage of reads mapping to the fungal transcriptome in each sample significantly correlated with measured lesion size (Supplementary Figure 5). Hierarchical clustering of the normalised expression data demonstrates high similarity between the biological replicates of each accession, with *B. cinerea* and *S. sclerotiorum* inoculated samples of an accession often also clustering together (Supplementary Figure 6). There is no clear grouping of the expression data by lettuce type.

Spearman correlations were calculated between lesion size and lettuce gene expression after both *B. cinerea* and *S. sclerotiorum* infection for each gene detected in our transcriptome profiling. After false discovery rate correction, 1,605 and 9,936 lettuce genes exhibited expression levels significantly correlated with *B. cinerea* and *S. sclerotiorum* lesion size, respectively (Supplementary Dataset 5). The difference in the number of genes showing correlation between expression and lesion size for each pathogen infection is likely due to the timing of sampling. Disease symptoms appear much faster following *B. cinerea* inoculation compared to *S. sclerotiorum*; hence, these samples are at a later stage of infection and the profiling has perhaps missed the critical dynamic period of transcriptome reprogramming. These genes with expression correlated to lesion size are likely to include many genes where the difference in expression between accessions is simply due to the dynamics of infection progression, rather than differences in gene expression being a potential driver of varying lesion size. For example, a gene upregulated during pathogen infection would likely have higher expression in a more susceptible accession simply because the infection has progressed faster and more tissue is responding to the pathogen. In contrast, genes downregulated during pathogen infection could have higher expression in a more resistant accession (compared to a susceptible accession) simply because infection has progressed more slowly and less plant tissue has responded. We therefore removed these categories of genes (upregulated genes correlated with susceptibility and downregulated genes correlated with resistance). To determine whether genes were up or downregulated during pathogen infection of lettuce, we used an RNAseq dataset comparing lettuce gene expression in leaves after *B. cinerea* or *S. sclerotiorum* inoculation to mock inoculation. Three biological replicates were harvested from leaves of the lettuce variety Saladin at 24 hpi with *B. cinerea* (and mock) and 42 hpi with *S. sclerotiorum* (and mock). A total of 8,130 (4,165 up/3,965 down) and 5,466 (3,329 up/2,137 down) genes were significantly differentially expressed in response to *B. cinerea* and *S. sclerotiorum*, respectively (Supplementary Dataset 6). Integrating this data with the diversity set RNAseq data and removing upregulated genes correlated with susceptibility and downregulated genes correlated with resistance resulted in 305 and 3,724 lettuce genes correlated with resistance to *B. cinerea* and *S. sclerotiorum*, respectively, as well as 326 and 1,580 correlated with susceptibility across the different accessions (Supplementary Dataset 5c, d). Of these, 174 genes correlated with resistance to both pathogens and 211 with susceptibility to both pathogens. Figure 3a illustrates the expression profiles for the 50 lettuce genes with the highest correlation with resistance against *S. sclerotiorum*, and Figure 3b shows the 50 genes with the highest correlation with susceptibility to *S. sclerotiorum*.

**Figure 3:**
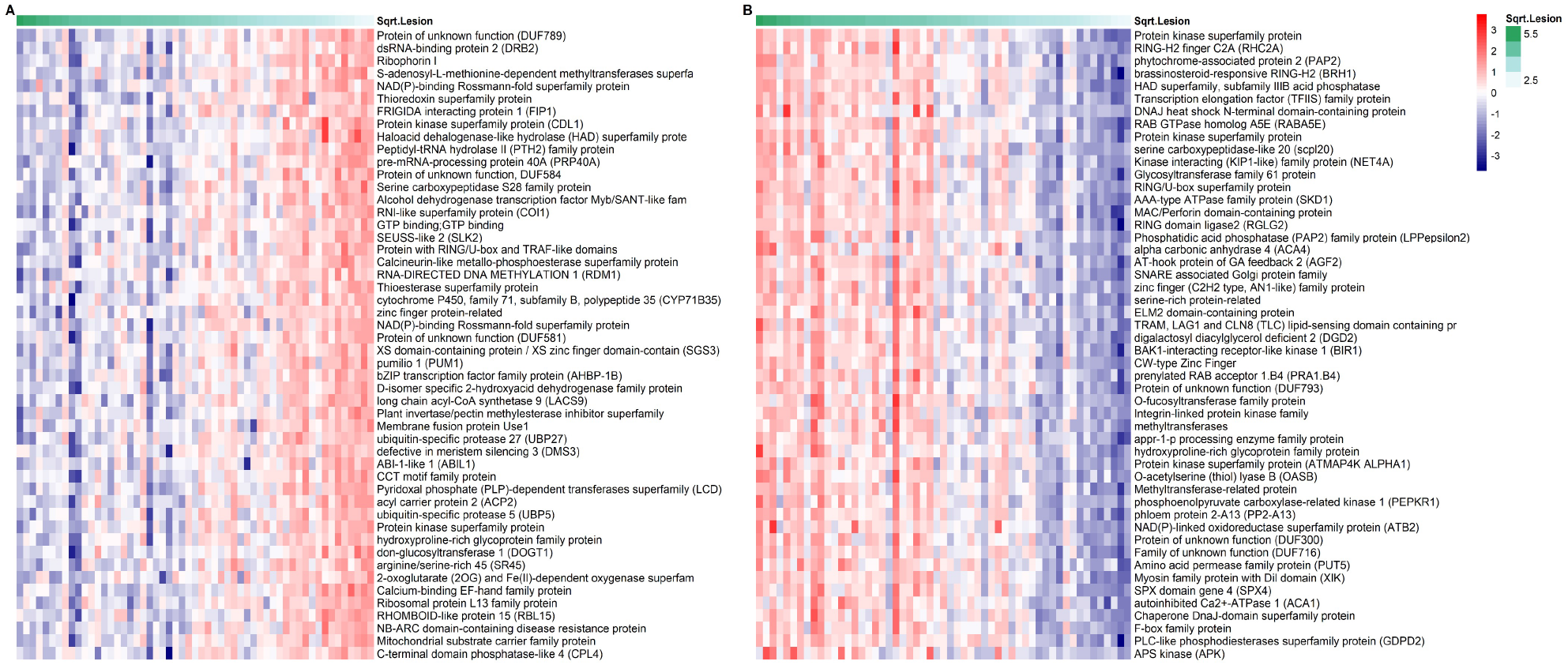
Heatmap of the top 50 lettuce genes identified with expression that correlates with *S. sclerotiorum* (A) resistance and (B) susceptibility across the diversity set accessions. The samples in the heatmap are ordered, with most susceptible (largest *S. sclerotiorum* lesion size) on the left and most resistant (smallest *S. sclerotiorum* lesion size) on the right.

The filtered lists of genes with expression significantly correlated with resistance or susceptibility contain several genes whose Arabidopsis orthologs have a known role in defence against *B. cinerea* and *S. sclerotiorum*, providing an initial validation of the data and indicating the ability of our approach to identify genes acting both positively and negatively on host immunity against these pathogens. For example, expression of two lettuce orthologs (Lsat_1_v5_gn_2_122000 and Lsat_1_v5_gn_9_61461) of coronatine insensitive 1 (COI1), the jasmonic acid receptor that is required for defence against *B. cinerea* (Feys et al. 1994; Rowe et al. 2010), are significantly inversely correlated with *S. sclerotiorum* lesion size (RS = -0.67 and -0.61). Two orthologs of TOPLESS (TPL) (Lsat_1_v5_gn_1_31280 and Lsat_1_v5_gn_5_63700) are significantly correlated with *S. sclerotiorum* resistance (R= -0.59 and -0.57, respectively). Arabidopsis triple mutants of TPL and the highly similar TOPLESS-related proteins (TPRs) 1 and 4, *tpl1*/*tpr1/tpr4*, show increased susceptibility to *B. cinerea* (Harvey et al. 2020). In addition, an ortholog of MAP kinase substrate 1 (MKS1), Lsat_1_v5_gn_1_8801, has expression correlated with *S. sclerotiorum* susceptibility (R=0.65) and in Arabidopsis, MSK1 is known to directly bind the key defence regulator WRKY33 with overexpression of MKS1 resulting in *B. cinerea* susceptibility (Petersen et al. 2010). These examples demonstrate our ability to identify defence genes from this dataset and increase our confidence in identifying novel lettuce defence components.

### GO-term analysis reveals enrichment of lettuce RNA binding proteins amongst genes with expression correlated with S. sclerotiorum resistance

We examined the biological processes represented by genes correlated with the defence response across the diverse lettuce accessions using GO-term enrichment. Lettuce genes have poor GO annotation; therefore, we performed GO-term enrichment analysis using the Arabidopsis orthologs of the lettuce genes correlated with resistance and susceptibility to *S. sclerotiorum* (2,985 and 1,254 genes respectively)(Figure 4, Supplementary Dataset 7). Strikingly, amongst the GO-terms enriched in genes correlated with *S. sclerotiorum* resistance were multiple terms associated with post-transcriptional RNA processing and regulation including gene silencing (GO:0016458), RNA interference (GO:0016246), dsRNA processing (GO:0031050), RNA modification (GO:0009451), exosome RNase complex (GO:0000178), and RNA splicing (GO: GO:0008380). Genes correlated with increased susceptibility to *S. sclerotiorum* were enriched for vesicle transport and cell growth related GO-terms.

**Figure 4:**
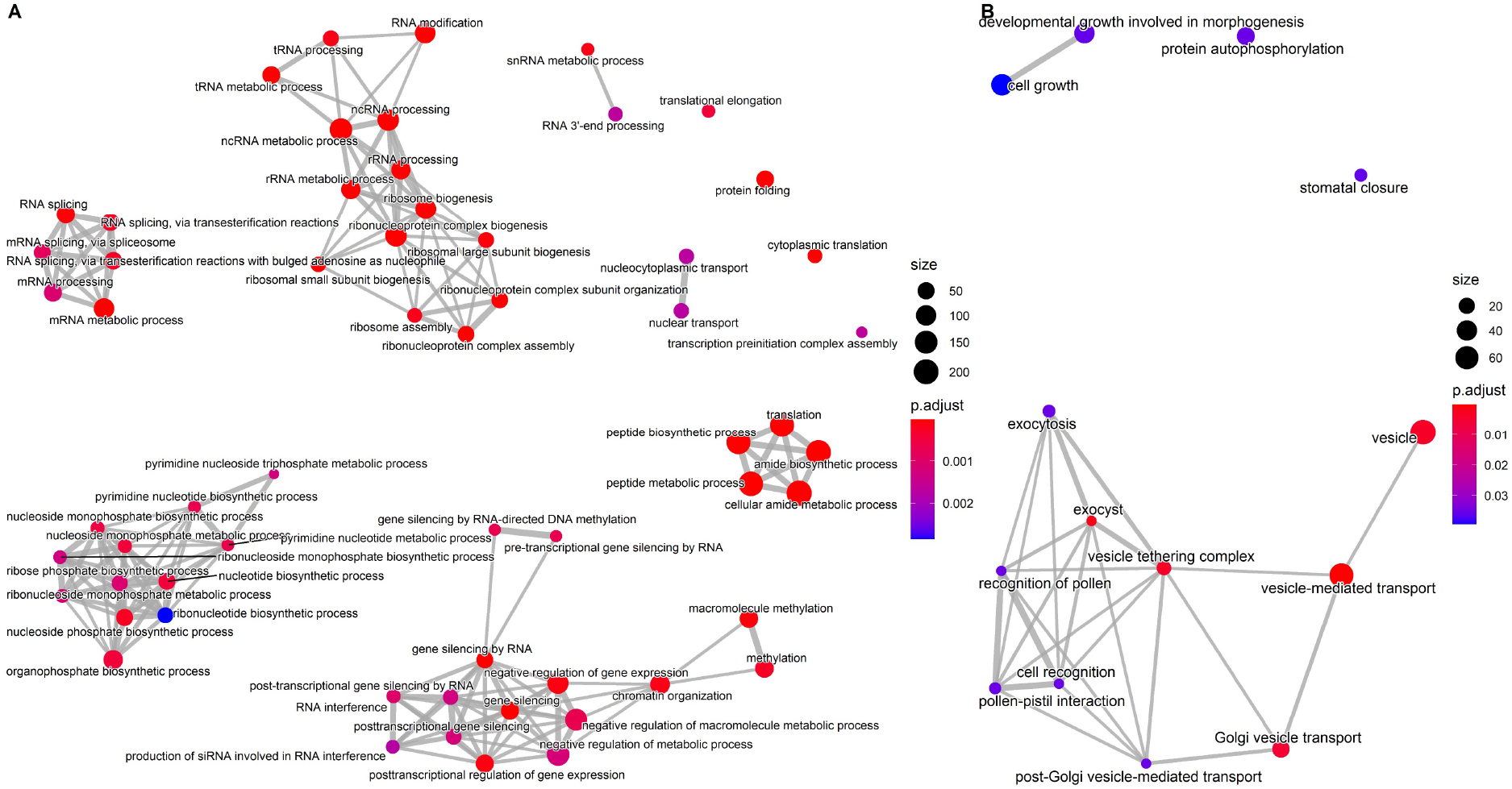
Gene Ontology (GO)-term enrichment networks of Arabidopsis orthologs of *S. sclerotiorum* (A) resistance and (B) susceptibility correlated genes. Each node is a statistically enriched GO-term (against background of all Arabidopsis genes with an identified lettuce ortholog). Node colour represents the relative level of statistical significance of the GO-term. Edges represent GO-terms with overlapping genes.

As shown above, *S. sclerotiorum* resistance correlated genes show a remarkable enrichment for GO-terms involved in RNA production, processing and RNA-mediated regulation. Post-transcriptional gene regulation via small RNAs is known to be a critical component of the host immune response and to contribute to reciprocal host–pathogen manipulation during infection by different types of plant pathogens (Huang et al. 2019). Seventy-two lettuce genes significantly correlated with resistance to *S. sclerotiorum* were identified as orthologs of Arabidopsis genes involved in gene silencing (GO:0016458)(Supplementary Dataset 7c). These 72 genes include several core components of the RNAi-mediated gene silencing pathway (Borges and Martienssen 2015) such as Dicer-like (DCL)2, DCL3, DCL4, Argonaute 1 (AGO1) and RNA-dependent RNA polymerase 2 (RDR2). In Arabidopsis, the gene silencing mutants *dcl4-2, ago9-1, rdr1-1, rdr6-11* and *rdr6-15* have been shown to increase susceptibility to *S. sclerotiorum* (Cao et al. 2016) while *dcl1* increased susceptibility to *B. cinerea* (Weiberg et al. 2013). Our data suggest a similar role for gene silencing in the defence response of lettuce against these broad host range pathogens.

Pentatricopeptide Repeat (PPRs) proteins are a family of RNA-binding proteins expanded in plants and involved in base editing and processing of organellar RNAs (Barkan and Small 2014), and their transcripts are a major source of secondary small interfering RNAs (siRNAs) in plants (Howell et al. 2007). The lettuce genome contains 513 putative PPRs (Reyes-Chin-Wo et al. 2017), 184 of which show correlation of expression with lesion size in our data. Expression of 178 lettuce PPRs is correlated with increased resistance to *S. sclerotiorum* (and six with increased susceptibility)(Supplementary Dataset 5) suggesting a potential key role for PPRs in the lettuce immune response. Figure 5 shows the expression profile of the top 20 resistance correlated PPRs.

**Figure 5:**
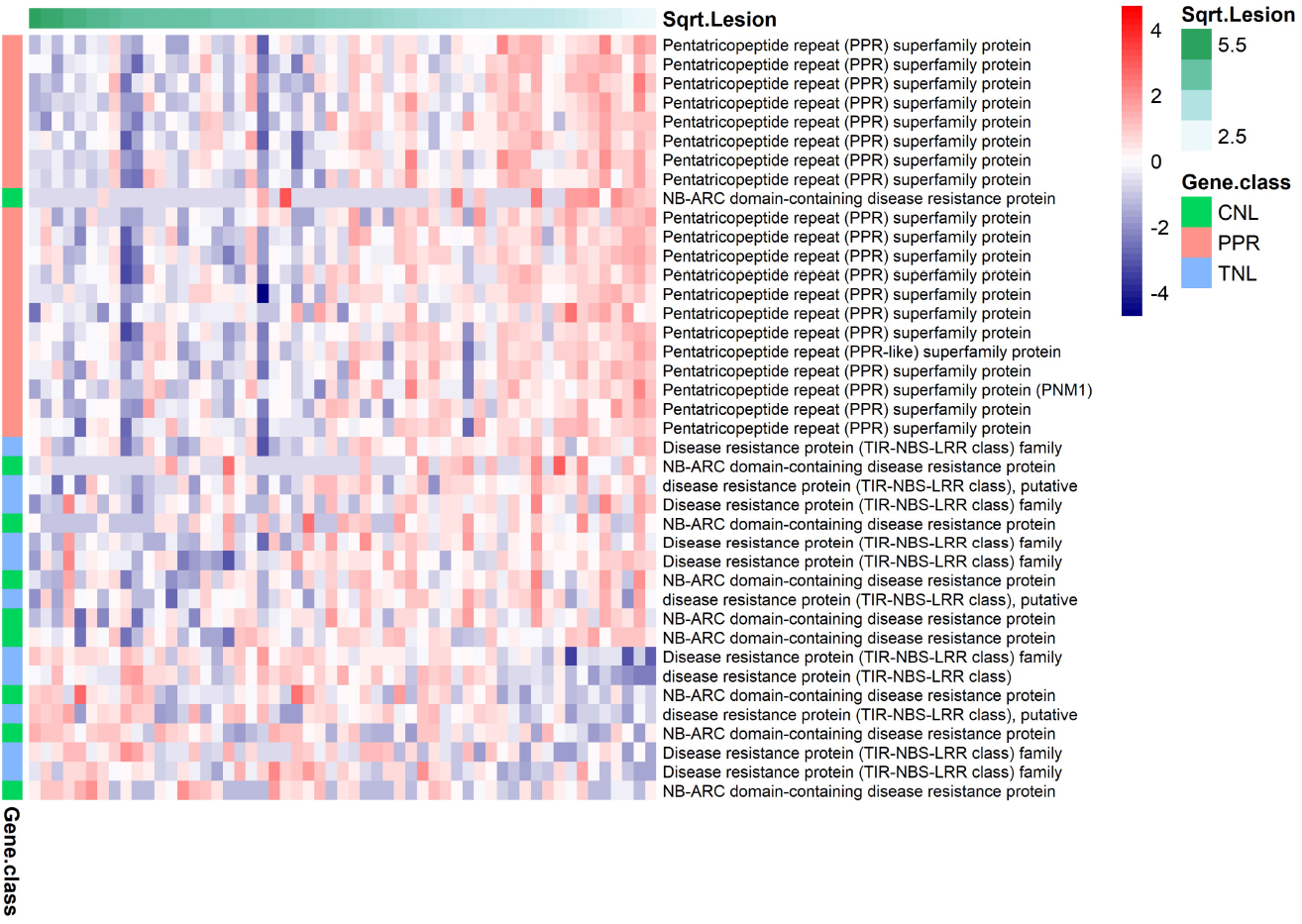
Expression of the top 20 pentatricopeptide repeat (PPR) genes whose expression is correlated with resistance against *S. sclerotiorum* (i.e. reduced lesion size) and all 20 nucleotide binding leucine-rich repeat (NLR) genes with expression correlated with *S. sclerotiorum* lesion size (12 correlated with resistance and eight with susceptibility). The NLRs are classified as Coiled-coil (CC)-NLRs (CNLs) or Toll-interleukin-1 receptor (TIR)-NLRs (TNLs). The individual lettuce samples are ordered left to right on the basis of lesion size after inoculation with *S. sclerotiorum*, with the most susceptible (largest lesion size) on the left and most resistant (smallest lesion size) on the right. Log_2_ expression is indicated by the red/blue scale.

### Multiple pathogen recognition genes have expression correlated with S. sclerotiorum lesion size across diverse lettuce accessions

Pathogen recognition by the host plant is mediated by both cell-surface (receptor-like kinases, RLKs and receptor-like proteins, RLPs) and intracellular (nucleotide binding site leucine-rich repeat proteins, NLRs) proteins. Our analysis suggests both groups of receptors impact lettuce resistance to *S. sclerotiorum*. Twenty-six RLKs and 14 RLPs had expression correlated with *S. sclerotiorum* lesion size; 12 RLKs and three RLPs with resistance and 14 RLKs and 11 RLPs with susceptibility (Supplementary Figure 7). Notably RLKs whose Arabidopsis orthologs have well-established and interacting roles in necrotrophic pathogen recognition were identified in this analysis. For example, mutants of BAK1 (BRI1-associated receptor kinase 1) show increased susceptibility to *B. cinerea* (Kemmerling et al. 2007) and expression of LsBAK1 (Lsat_1_v5_gn_9_117621) correlates with increased *S. sclerotiorum* resistance (R=-0.38). BAK1 directly interacts with the flagellin-sensitive receptor FLS2 (following flg22 perception) initiating downstream defence responses (Chinchilla et al. 2007) and a lettuce ortholog of FLS2, Lsat_1_v5_gn_7_32940, also had expression correlated with *S. sclerotiorum* resistance (R=-0.47). In contrast, BIR1 (BAK1-interacting receptor-like kinase 1), directly interacts with BAK1 (Ma et al. 2017) and negatively regulates defence (Gao et al. 2009). Consistent with this function, the expression of Lsat_1_v5_gn_0_1380, an ortholog of BIR1, was found to correlate with lettuce susceptibility to *S. sclerotiorum* (R=0.69). Our detection of this group of interacting known immune regulators provides confidence in our approach and suggests other RLKs identified could have genuine impacts on *S. sclerotiorum* and/or *B. cinerea* resistance. Similarly, there is a precedent for the involvement of RLPs in plant response to necrotrophic fungal infection with Arabidopsis RLP23 required for proper defence against *B. cinerea* (Ono et al. 2020).

In total, 236 genes in the lettuce genome encode intracellular NLRs, with 47 encoding coiled-coil NLRs (CNLs) and 187 encoding Toll/Interleukin-1 type NLRs (TNLs)(Christopoulou et al., 2015). Of these NLRs, 20 showed significant correlation of expression with lesion size after *S. sclerotiorum* infection with expression of 12 correlating with increased resistance and eight with increased susceptibility (Figure 5). As for the majority of the genes with expression correlated with lesion size, all 20 do not change in expression in response to pathogen infection (at least from our single time point dataset), suggesting that it is basal expression levels of these NLR genes that is impacting quantitative disease resistance to *S. sclerotiorum* in lettuce.

### Parents of existing lettuce mapping populations show quantitative variation in susceptibility to necrotrophic fungal pathogens

We screened the parents of existing lettuce mapping populations to test whether these populations would be suitable for dissecting the mechanistic basis of quantitative variation in resistance to *B. cinerea* and *S. sclerotiorum*. Seventeen lettuce accessions, the parents of 11 different mapping populations, were phenotyped using the detached leaf inoculation assay with both *B. cinerea* and *S. sclerotiorum*. Of the 11 parental combinations, two exhibited significantly different lesion size after *B. cinerea* inoculation, and five exhibited significantly different lesion size after *S. sclerotiorum* inoculation (Figure 6). The parents of two mapping populations exhibited significantly different lesion sizes in response to both pathogens. Notably, in the Greenlake x Diana cross, *S. sclerotiorum* lesions on lettuce variety Greenlake were significantly larger than lesions on variety Diana, while *B. cinerea* lesions were significantly smaller, indicating that Greenlake and Diana exhibit contrasting susceptibility to the two pathogens. The second set of parental lines, *L. sativa* PI251246 (Subbarao, 1998) and an Armenian *L. serriola*, showed consistent responses to *B. cinerea* and *S. sclerotiorum* with lesions caused by both pathogens larger on PI251246 than on leaves of the Armenian *L. serriola* line. This mapping population was investigated further to identify genomic regions mediating this difference in lesion development.

**Figure 6:**
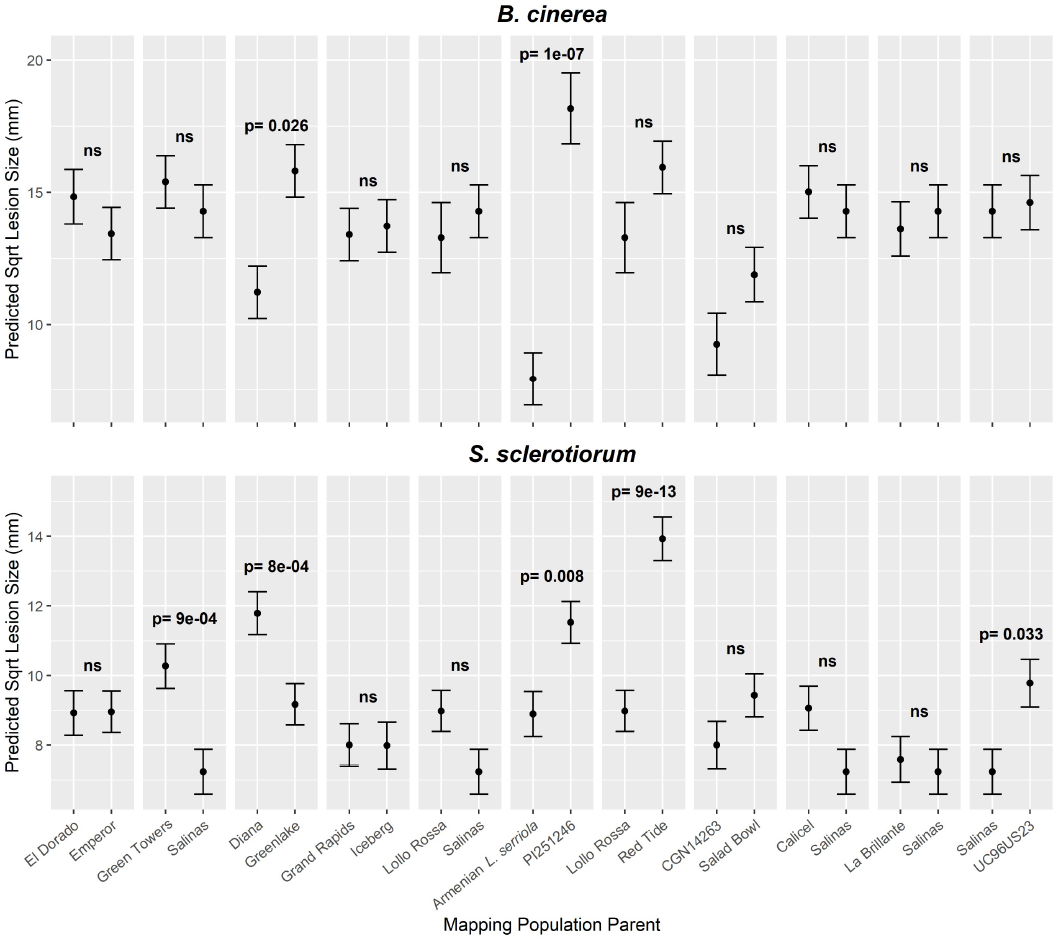
Parent lines of lettuce mapping populations differ in lesion size after inoculation with *B. cinerea* or *S. sclerotiorum*. REML predicted square root mean lesion size of *B. cinerea* (top) or *S. sclerotiorum* (bottom) on detached lettuce leaves of mapping population parents available from UC Davis. Lines are shown grouped as parents of mapping populations. Multiple cases of the same line represent one set of data that is repeated to allow comparison within a different parental pair. Error bars are REML predicted standard error, where n is between 15 and 29. Tukey HSD p-values are shown where there is significant difference, otherwise ‘ns’ indicates not significant.

### Five genomic regions mediating lettuce resistance to fungal pathogens in a detached leaf assay

A total of 234 F6 recombinant inbred lines (RILs) resulting from the cross between the Armenian *L. serriola* and PI251246 (*L*. sativa) were phenotyped in a replicated incomplete experimental design using both *B. cinerea* and *S. sclerotiorum* detached leaf assays (Supplementary Dataset 8a). QTL mapping identified five loci impacting lesion size following inoculation with *B. cinerea* or *S. sclerotiorum* (Figure 7, Supplementary Figure 8). Three QTL impacted the size of *S. sclerotiorum* lesions and two impacted the size of *B. cinerea* lesions. Despite correlation between resistance to these two pathogens across the lettuce diversity set (Supplementary Figure 2) and the parental lines demonstrating significant differences in response to both pathogens (Figure 6), QTL mediating quantitative resistance to the two pathogens did not co-locate. This suggests five QTL exist between these two parent lines independently contributing to disease resistance. No evidence for epistatic interactions between the QTL loci was detected.

**Figure 7:**
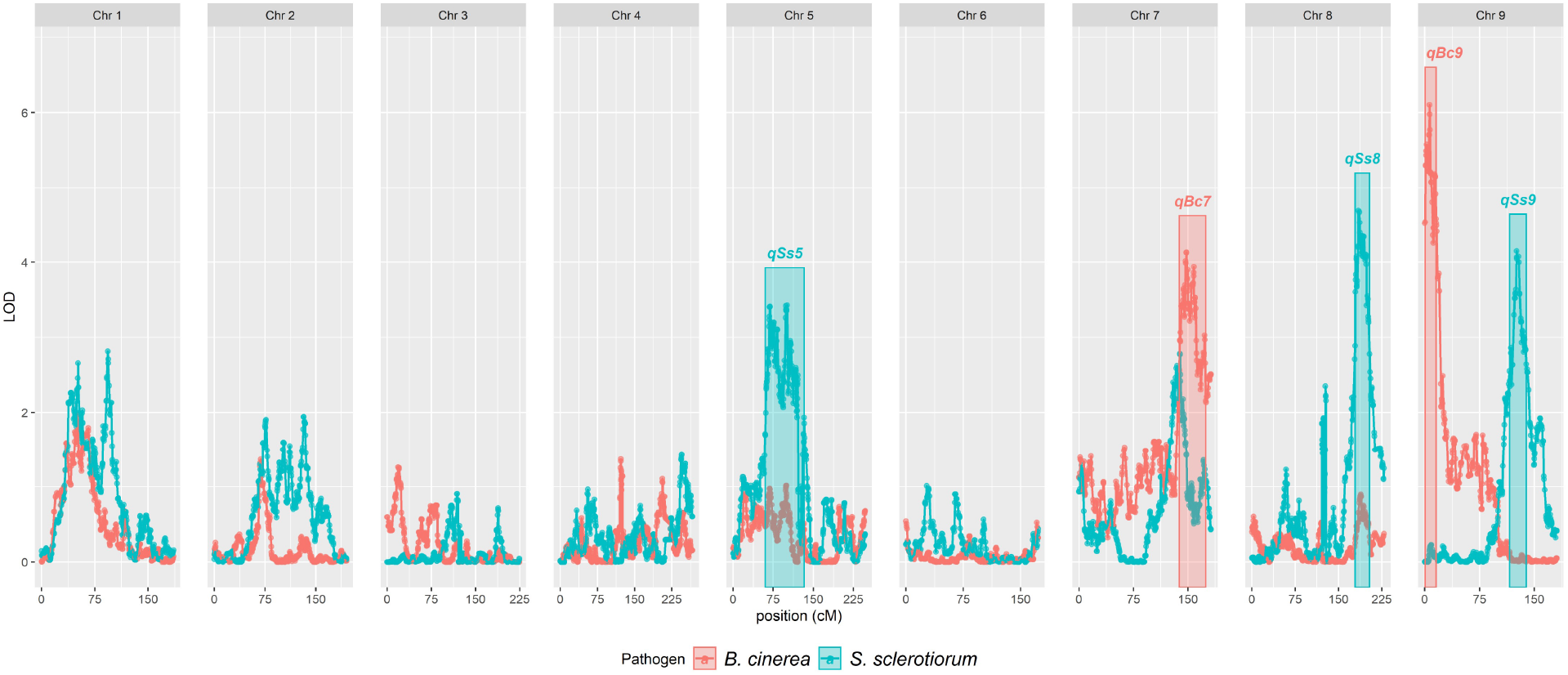
Quantitative trait loci associated with reduced lesion size of *B. cinerea* or *S. sclerotiorum*. LOD scores from ’stepwiseqtl’ multi-QTL selection models using the Haley-Knott algorithm, genotyping-by-sequencing markers and predicted mean lesion size from the detached leaf assay data for each pathogen are shown. Data relating to *B. cinerea* inoculation are shown in red, whereas those from *S. sclerotiorum* inoculation are shown in blue. Five significant QTL (*qSs5,qSs8, qSs9, qBc7* & *qBc9*) were maintained in the final model after backwards elimination of insignificant loci. Boxes represent the 1.5LOD confidence intervals around the peak LOD of each QTL. The nine lettuce chromosomes are shown along the x-axis.

Each QTL explains between 7 and 11% of the lesion size variation (Table 1). For four of the five QTL, the alleles conferring reduced lesion size were derived from the Armenian *L. serriola* parent, which showed increased resistance to *B. cinerea* and *S. sclerotiorum* compared to the other parental line, PI251246. However, at *qSs9* (*S. sclerotiorum* QTL on Chromosome 9), the resistance allele originates from the susceptible parent, PI251246. This suggests the presence of alleles with both positive and negative effects on disease resistance, which can be separated by recombination. RILs that contain the resistance alleles at both *B. cinerea* QTL or all three *S. sclerotiorum* QTL have significantly reduced lesion size compared to RILs with susceptibility alleles at the same loci (Supplementary Figure 9).

**Table 1:** Summary statistics for resistance quantitative trait loci (QTL) identified in the Armenian *L. serriola* x PI251246 mapping population. The position along the chromosome, peak LOD score, variance in lesion size explained, genetic and physical size, and the number of genes present (and expressed in our data sets) is indicated for each QTL.

| QTL | Pathogen | Chr | Pos (cM) | pLOD | Variance Exp | Genetic size (cM) | Physical size (Mb) | #Genes | #Genes expressed |
| --- | --- | --- | --- | --- | --- | --- | --- | --- | --- |
| qBc7 | *B. cinerea* | 7 | 148.0 | 4.1 | 7.29% | 36.5 | 34.9 | 639 | 313 |
| qBc9 | *B. cinerea* | 9 | 6.4 | 6.1 | 10.75% | 15.0 | 26.7 | 776 | 415 |
| qSs5 | *S.sclerotiorum* | 5 | 101.0 | 3.4 | 7.02% | 72.7 | 118.4 | 1353 | 565 |
| qSs8 | *S.sclerotiorum* | 8 | 184.5 | 4.7 | 8.14% | 25.1 | 40.1 | 292 | 142 |
| qSs9 | *S.sclerotiorum* | 9 | 125.6 | 4.1 | 8.54% | 22.8 | 18.8 | 204 | 83 |

**Table 2.**
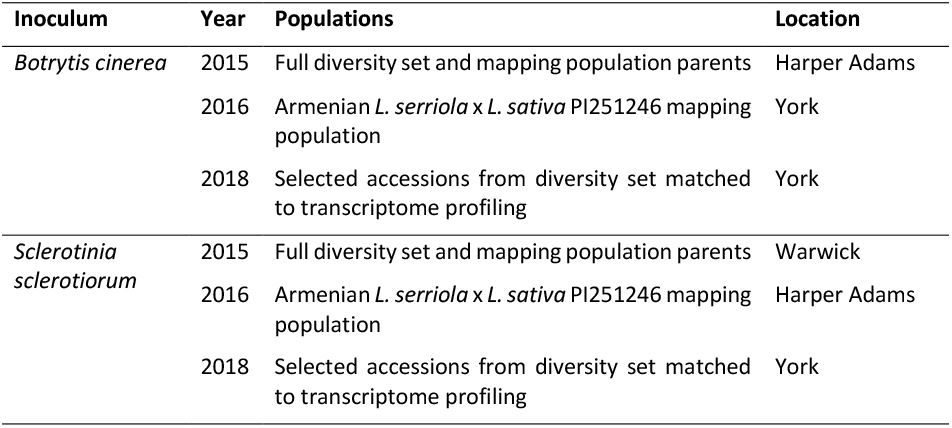
Location of detached leaf inoculation experiments

To define boundary markers for each QTL, confidence intervals of 1.5 LOD surrounding each QTL peak were calculated using ‘*lodint’*, which were expanded to the next marker. QTL boundary markers were mapped onto the *L. sativa* cv Salinas v8 genome (Reyes-Chin-Wo et al. 2017). Predicted genes positioned between the QTL boundary markers in the Salinas cultivar could then be identified (Table 1, Supplementary Dataset 8d). A large variation in QTL size (and gene number) was observed, with *qSs5* the largest at 72.7 Mb and containing 1,353 genes. The smallest QTL (by gene number) is *qSs9*, at 22.8 Mb with the region containing 204 genes.

### Identification of candidate casual genes in the QTL through transcriptome profiling

We attempted to predict causal genes within the identified QTL regions using transcriptome and genome data. Both parent lines, the Armenian *L. serriola* and PI251246, were included in the lettuce diversity set whose transcriptomes were profiled 48 hpi and 43 hpi with *B. cinerea* and *S. sclerotiorum*, respectively (Supplementary Dataset 3). We carried out a second inoculation of these parental lines and generated an additional four biological replicates of transcriptome profiles (48 hpi after *B. cinerea* and 64 hpi after *S. sclerotiorum* inoculation). Principal component analysis of the 28 samples (three/four replicates x two experiments x two pathogens x two lettuce lines) demonstrates clear differences between the parental lines and similarity of the replicates within and across experiments (Supplementary Figure 10). Prior to any differential expression analysis, numbers of potential QTL candidate genes could be reduced by 46-59%, by removing genes which failed to pass the low-expression filter, i.e they had no detectable expression after either *B. cinerea* or *S. sclerotiorum* infection (Table 1). Differential expression analysis was carried out on each experiment/pathogen combination separately with a minimum ±1.2 log2 fold change threshold to filter significantly differentially expressed genes (DEGs).DEGs and their expression values are available in Supplementary Dataset 9. In response to *S. sclerotiorum* inoculation, there were 1,198 DEGs (425 up/773 down) across both sets of data with 96 (24 up/72 down) of these DEGs located within identified resistance QTL. One hundred forty-five DEGs (58 up/87 down) were common to both *B. cinerea* inoculation datasets with 10 (5 up/5 down) of these located within QTL. Ninety genes (36 up/54 down) were differentially expressed (in the same direction) between the parent lines in response to both pathogens, of which seven (three up/four down) are in QTL.

For the lettuce genes located within the QTL regions, we integrated the parental line transcriptome data above with information on whether expression of the gene in the lettuce diversity set was correlated with lesion size in response to pathogen infection to predict candidate causal genes underlying the identified resistance QTL (Figure 8). This analysis highlighted a number of genes with expression patterns consistent with a potential role in mediating fungal pathogen resistance within this mapping population; for example, Lsat_1_v5_gn_5_91640 (LsPDR12) is an ortholog of the Arabidopsis gene Pleiotropic Drug Resistance 12 (AtPDR12), known to mediate camalexin secretion in response to *B. cinerea* infection (He et al. 2019). LsPDR12 is significantly correlated with *S. sclerotiorum* resistance in the diversity set (RS = -0.37), is upregulated in Armenian *L. serriola* compared to PI251246 in both experiments and is located within the QTL *qSs5* (for which the resistance allele comes from the Armenian *L. serriola* parent). Another potential candidate, Lsat_1_v5_gn_9_1180, encodes an ortholog of CML38, is located within the *qBc9* QTL and is upregulated in PI251246 compared to the Armenian *L. serriola*, i.e. higher expression in the susceptible parent. However, consistent with this, Arabidopsis CML38 promotes degradation of suppressor of gene silencing 3 (SGS3)(Field et al. 2021), the lettuce ortholog of which is correlated with resistance against *S. sclerotiorum* (Supplementary Dataset 5).

**Figure 8:**
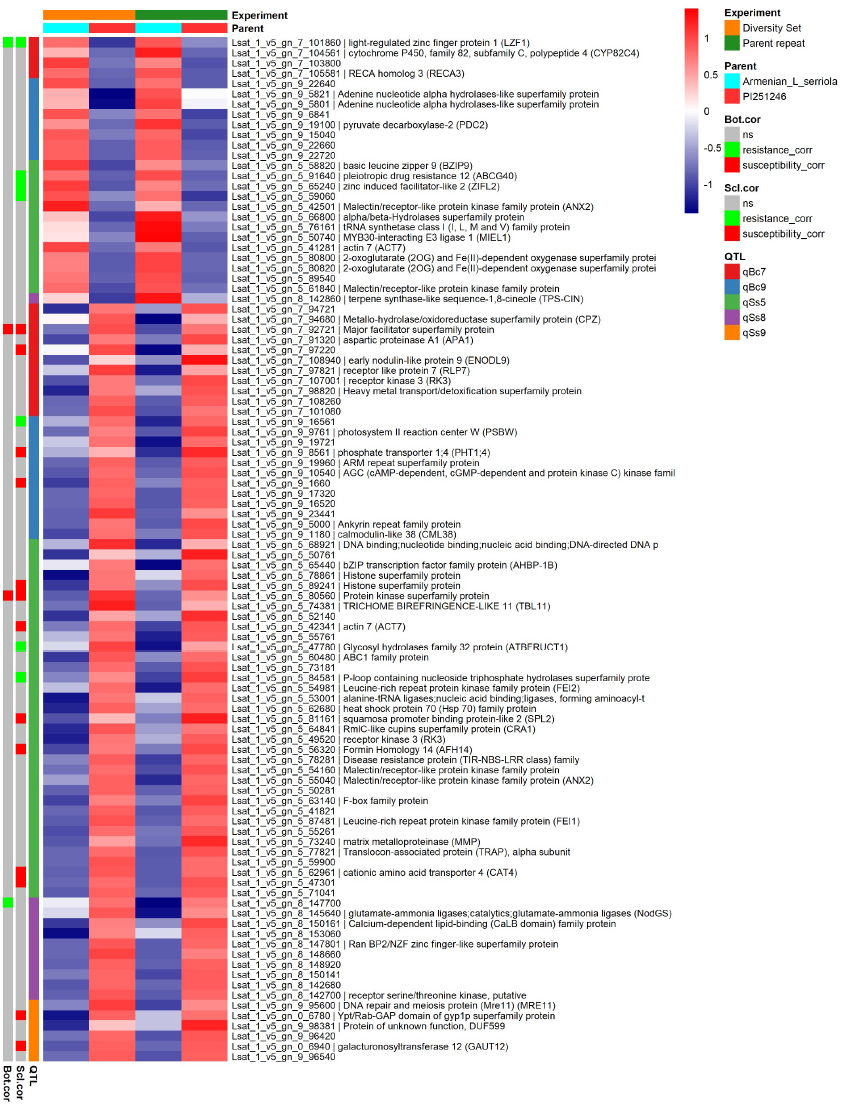
Integration of gene expression data with QTL to predict potential causal genes. The expression of genes differentially expressed between the mapping population parents (Armenian *L. serriola*, PI251246,) after *S. sclerotiorum* infection in two datasets (as part of the diversity set and a specific repeat of the two lines) and located within a QTL are shown. For all QTL, except for qSs9, the resistance allele originates in the Armenian *L. serriola* line. The two columns on the left indicate genes whose expression is correlated with pathogen resistance or susceptibility in the lettuce diversity set (ns = no significant correlation).

## Discussion

We demonstrated that genetic variation for resistance to the fungal pathogens *S. sclerotiorum* and *B. cinerea* exists within a lettuce diversity set comprised of both *L. sativa* varieties and wildtype relatives (Figure 1). Quantitative resistance within this diversity set was assessed using a detached leaf assay. Although this does not necessarily correlate with field resistance, it has the advantage of reliable and consistent inoculation, and of measuring immunity in a manner that is not dependent on the overall plant architecture. In this way, we hope to identify traits that could be exploited in a range of lettuce morphotypes. Crucially, we used inocula of spore suspensions for both pathogens, which mimics the natural infection route, whereas most publications investigating variation in resistance against *S. sclerotiorum* use mycelial plugs (e.g. (Chittem et al. 2020) due to the difficulties, and time, taken to produce ascospores.

We identified parents of existing lettuce mapping populations that differed in susceptibility to the two fungal pathogens (Figure 6) and five QTL that impacted quantitative resistance differences between *L. sativa* PI251246 and an Armenian *L. serriola* (Figure 7). For four of these QTL the resistance allele originated in the Armenian *L. serriola* line. Although PI251246 was the more susceptible parent in this work, this accession was previously shown to have lower *S. sclerotiorum* disease incidence, but similar disease severity to a standard lettuce Butterhead variety, Rachel, in an inoculated glasshouse trial (Whipps et al. 2002). This suggests that PI251246 is able to escape *S. sclerotiorum* (due to architecture and/or rapid bolting) but lacks tissue resistance once infection becomes established. This is consistent with our results from detached leaf assays and a field trial where observed resistance of PI251246 to *Sclerotinia minor* was attributed to rapid bolting characteristics (Hayes et al. 2010). These architectural and rapid bolting attributes would not be beneficial in cultivated lettuce. In contrast, the Armenian *L. serriola* had significantly higher resistance than PI251246 to both S. *sclerotiorum* and *B. cinerea* in the detached leaf assay, suggesting this accession may have beneficial traits which could be exploited in lettuce varieties with varying architectures. *L. serriola* is a wild lettuce believed to be the progenitor of domesticated *L. sativa* (Uwimana et al. 2012). Wild relatives of crop plants are often sources of disease resistance and in previous work with this mapping population, four QTL conferring resistance to *Verticillium dahliae* have been identified (Sandoya et al. 2021) with all four beneficial alleles from the Armenian *L. serriola* parent.

Although QTL have been identified for field resistance to *S. minor* (Mamo et al. 2019), to our knowledge these are the first lettuce QTL reported for resistance against *S. sclerotiorum* and *B. cinerea*. Multiple *S. sclerotinia* resistance QTL mapping studies exist in sunflower and *Brassica napus* (for example, (Behla et al. 2017), as well as a small number of GWAS in soybean and *B. napus*, all excellently reviewed in (Wang et al. 2019). As in our study, where the QTL explained between 5% and 10% of the variation, published QTL have minor effects on disease resistance (less than 10%). A similar polygenic basis for resistance has been seen against *B. cinerea* with QTL studies in Arabidopsis, *Solanum* species and *Brassica rapa* and GWAS in Arabidopsis (Corwin and Kliebenstein 2017).

Although the *S. sclerotinia* isolate used here was initially obtained from field grown lettuce, our analysis was restricted to single isolates of both pathogens. Previous studies using *B. cinerea* have indicated that there is a high level of isolate-specificity to quantitative resistance loci (Zhang et al. 2016) and data on disease outcome using 98 *B. cinerea* isolates and 90 genotypes of eight plant hosts (including lettuce) demonstrated a much greater impact on lesion size from the *B. cinerea* isolate (40-71%) than the host genotype (3-8%)(Caseys et al. 2021). Similarly, in *B. napus* both pathotype-specific and pathotype-independent resistance against *S. sclerotiorum* has been identified (Neik et al. 2017). Hence, determining whether the QTL identified here can mediate resistance to a broad range of pathogen isolates will be critical to their value in crop improvement.

Despite multiple QTL/GWAS analyses, relatively little is known about the molecular mechanisms driving resistance, and the small effect of each QTL increases the complexity of the fine-mapping process. Hence, we used transcriptome data to predict potential causal genes. We go beyond just simply comparing differential expression in two contrasting parental lines (e.g. (Qasim et al. 2020) or integrating expression and genetic location of DEGs (Zhao et al. 2015), to identifying genes whose expression is correlated with resistance or susceptibility (as judged by lesion size) across 21 different lettuce accessions (Figure 3). Although this correlation analysis (unlike expression QTL analysis) does not pinpoint the genetic region responsible for variation in expression, it does enable us to identify genes that impact infection outcome (positively or negatively) but are not necessarily differentially expressed during infection. Furthermore, the use of multiple accessions (rather than the commonly seen comparison of one resistant and one susceptible line) gives us better ability to identify expression differences genuinely contributing to disease resistance. This analysis identified genes whose increase in expression is correlated with smaller lesion size (resistance) and larger lesion size (susceptibility).

One key group of genes that we identified as highly correlated with disease resistance were those involved in RNA-mediated post transcriptional regulation, in particular gene silencing. In well-studied plants, such as Arabidopsis and tomato, gene silencing (via microRNAs, miRNAs, or small interfering RNAs, siRNAs) is known to play a crucial role in the host immune response - both in regulating expression of plant genes, as well as silencing of genes in the pathogen (Qiao et al. 2021). Our data suggest a similar role for gene silencing in lettuce, and the prominence of orthologs of genes known to increase susceptibility in Arabidopsis to both *S. sclerotiorum* and *B. cinerea* provides confidence in this approach to highlight key mechanisms of quantitative resistance. One such potential mechanism is PPR-driven silencing of pathogen virulence genes. Increased expression of many lettuce *PPR* genes was correlated with decreased lesion size after pathogen inoculation (Figure 5). Intriguingly *PPR* transcripts are a major source of secondary siRNAs in plants, whose generation is triggered by both direct miRNA binding to specific *PPR* transcripts and via miRNA-mediated generation of trans-acting siRNAs (tasiRNAs)(Howell et al. 2007). Furthermore, siRNAs derived from *PRR* transcripts accumulate after *Phytophthora capsici* infection, can potentially target known pathogen virulence genes, and an effector from the pathogen can suppress accumulation of these siRNAs to promote infection (Hou et al. 2019). Arabidopsis RNA-dependent RNA polymerase 6 (RDR6) is required for generation of siRNAs from endogenous transcripts, and *rdr6* mutants show enhanced susceptibility to *B. cinerea* (Cai et al. 2018). Furthermore, mutation of the Pentatricopeptide repeat protein for germination on NaCl (PGN) in Arabidopsis led to increased susceptibility to *B. cinerea* (Laluk et al. 2011). Clearly, further research is needed to determine the importance and mechanism of *PPR* siRNA production in lettuce, especially given that in a comparative study across nine plant species, several new and potentially species-specific miRNAs were shown to drive production of these siRNAs (Xia et al. 2013).

Small RNAs are also thought to play a key role in regulating expression of NLR genes, helping regulate their expression (and hence inadvertent triggering of the defence response) in the absence of infection. NLRs are mostly known as an integral part of effector-triggered immunity (ETI), whereby pathogen effectors are directly or indirectly (as guards or decoys of effector targets) recognised by NLRs (Cui et al. 2013). As such, they have typically been associated with qualitative (all or nothing) disease resistance. Due to the lack of complete resistance phenotypes against broad host range necrotrophic fungal pathogens and very limited number of NLR genes shown to impact resistance, it was thought that NLR proteins and ETI are not important in defence against these pathogens (Mengiste 2012). However, in our data expression of multiple lettuce NLRs was shown to be correlated with resistance suggesting that, in addition to their well-known role in lettuce resistance against biotrophic pathogens (Simko et al. 2013; Parra et al. 2016, 2021), NLR genes in lettuce may play a role in quantitative resistance against *B. cinerea* and *S. sclerotiorum*. NLRs show huge diversity both within a single genome and in populations, and a multitude of incomplete NLRs (lacking one or more of the canonical domains but thought to still be able to function as adapters or helpers for other NLRs) are also found in all plants (Baggs et al. 2017). Several lettuce incomplete NLRs had expression correlated with *S. sclerotiorum* lesion size (Supplementary Figure 7) and there is a precedent for incomplete NLRs impacting resistance to broad host range necrotrophic pathogens, with mutations in the Arabidopsis gene *RLM3* (containing TIR and nucleotide binding domains) causing increased susceptibility to these pathogens, including *B. cinerea* (Staal et al. 2008). We also noted in our analysis that there were several NLRs whose increased expression was correlated to increased susceptibility to *S. sclerotiorum* (Figure 5) suggesting that NLRs may have both a positive and negative effect on resistance to this pathogen. Indeed, a Toll interleukin-1 receptor (TIR) type NLR in Arabidopsis, *LAZ5*, has been shown to increase susceptibility to *S. sclerotiorum* infection, with *laz5-1* mutants showing increased resistance (Barbacci et al. 2020).

Although the lettuce diversity set we used here is small compared to a collection that has been recently genome sequenced (Wei et al. 2021), our analysis demonstrated useful genetic variation for quantitative disease resistance, indicated crosses that could be useful in mapping this trait and identified multiple potential mechanisms for experimental testing. It is likely that different lettuce lines harbour different quantitative resistance mechanisms, and our gene expression correlation analysis has identified strong candidates for experimental testing that are not obviously segregating in the Armenian *L. serriola* x PI251246 population. However, combining transcriptome data from the parents and diversity set with QTL analysis has also identified a small number of potential causal genes for the resistance QTL in this population, with the strongest candidate being the lettuce ortholog of Arabidopsis Pleiotropic Drug Resistance 12 (AtPDR12) within the QTL *qSs5* (Figure 8). Obviously, the molecular mechanism underlying the resistance QTL may not necessarily be regulatory variation of a gene within the QTL itself (and hence identifiable in our analysis approach) but could be driven by sequence variation driving changes in post-transcriptional gene regulation or protein function.

In summary, we have identified multiple potential architecture-independent resistance mechanisms that may be successful for enhancing disease resistance in lettuce. Future work will aim to validate candidate genes, for example via fine-mapping of QTL and/or the generation of lettuce lines with gain or loss of function mutations/transgenes. The resistance traits could be incorporated into cultivated varieties (via marker-assisted selection) with genome editing of validated candidate genes offering an exciting route to exploit the genetic variation from these lettuce accessions without losing the beneficial traits stacked up in elite breeding lines.

## Author Contribution Statement

The study was conceived and designed by KD, JC, CY, FG, PH and DP. Experimental work and initial data analysis were carried out by AT, GH, JB, JG, AT, AJ and AG with further data analysis and data interpretation performed by HP, KD, JC, PH and DP. The mapping population was generated and genotyped by MT and RM. The manuscript was written by AT, HP and KD with input from all authors. All authors have approved the submission of this manuscript.

## Acknowledgements

We would like to thank the UK Vegetable Genebank housed at the University of Warwick for provision of seed of the lettuce diversity set and Sally James of the University of York Biology Technology Facility for generating the RNA sequencing libraries. We also thanks Josie Brough for technical assistance.

## Data Availability

All data is available either within the supporting information of this manuscript or in the NCBI Short Read archive under Bioproject PRJNA804213 (diversity set and mapping population parent RNAseq data) and Bioproject PRJNA808232 (single time point infected versus mock inoculated).

## Figure Legends

**Supplementary Figure 1:**
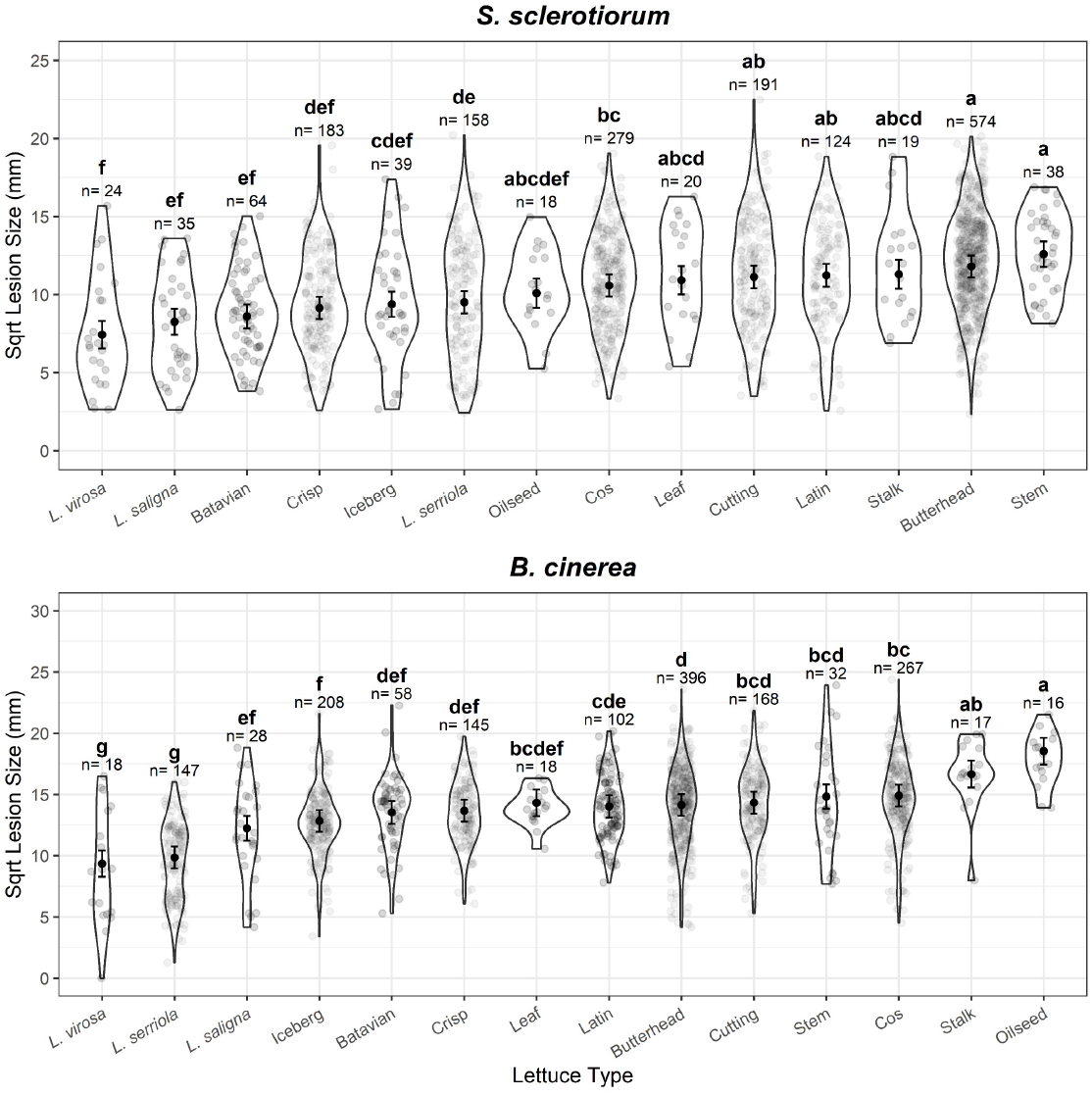
Variation in lesion size after inoculation with *B. cinerea* or *S. sclerotiorum* between lettuce types. Square root lesion size in response to *S. sclerotiorum* (top) or *B. cinerea* (bottom) on detached lettuce leaves of the Lettuce Diversity Fixed Foundation Set. Grey points show individual measured lesions, violins show the distribution. Black points show REML predicted (accounting for random variation between experimental replicates). Error bars indicate REML predicted standard error. Letters shown represent Tukey HSD significance groupings (p< 0.05). n = the number of lesions measured from each type in response to each pathogen.

**Supplementary Figure 2:**
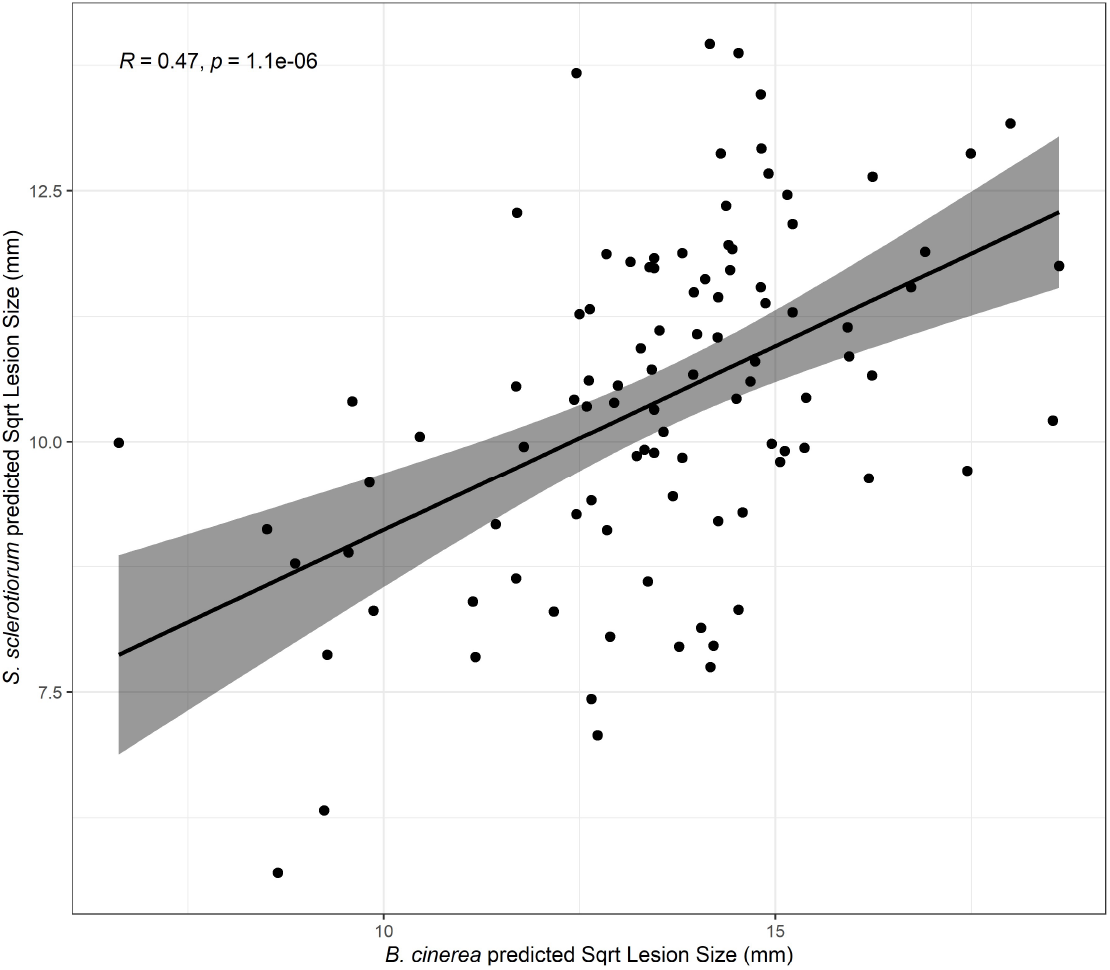
Correlation between lesion size 64 hours post inoculation with *S. sclerotiorum* and *B. cinerea*. Least-squares predicted mean square root lesion size of *B. cinerea* (x-axis) vs. *S. sclerotiorum* (y-axis) on detached lettuce leaves, where n ranges from two to 20 for each accession/pathogen combination. Linear regression line is shown in black, with 95% confidence intervals shaded in grey.

**Supplementary Figure 3:**
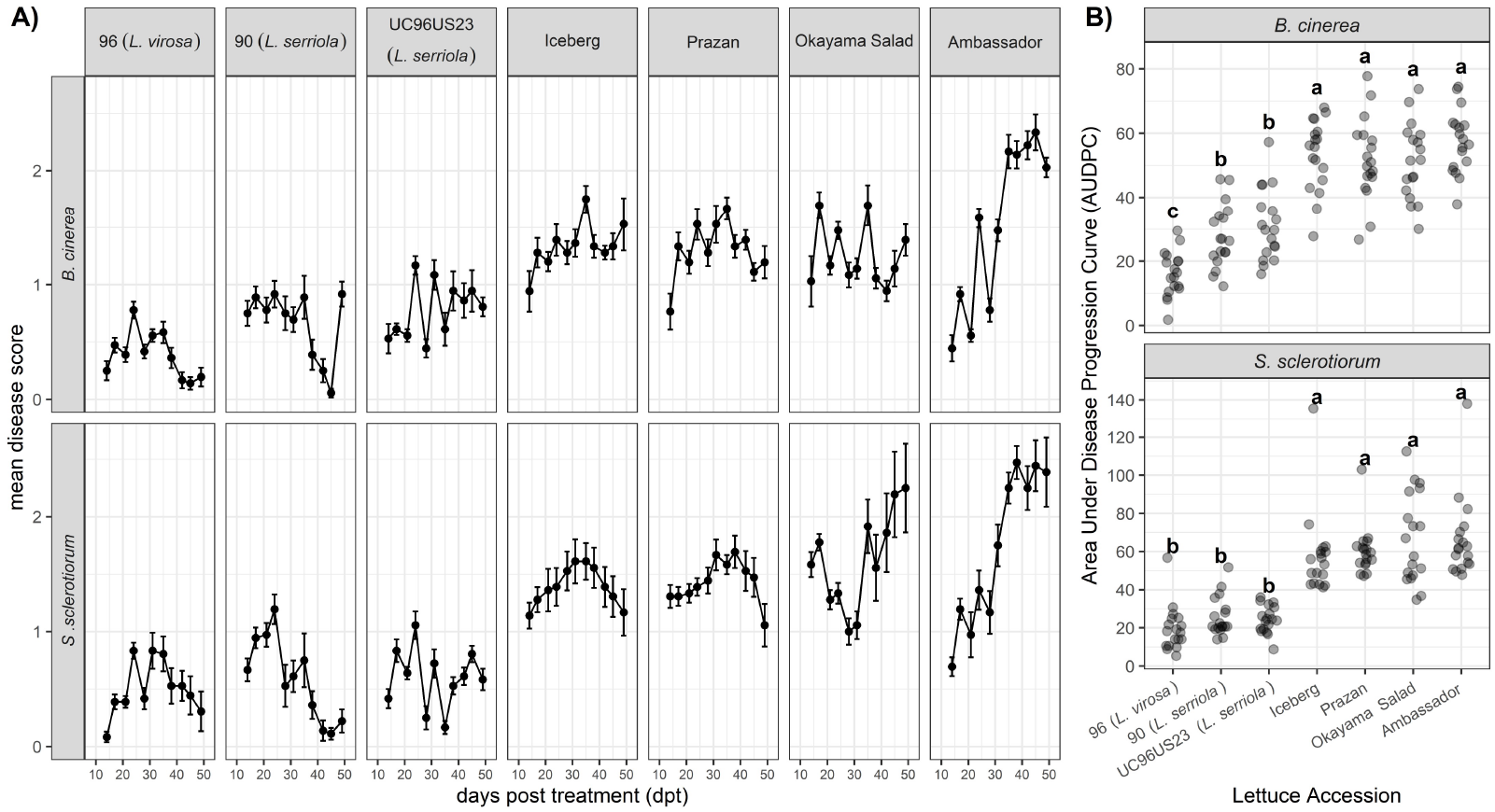
Whole Plant Disease incidence scoring on seven lettuce accessions. All accessions are *L. sativa* unless otherwise indicated. (A) Mean disease symptom scoring out of 4 for each lettuce accession from 14 to 49 days post treatment in response to *B. cinerea* and *S. sclerotiorum*. Error bars are standard error, n=18. (B) Area under the disease progression curve (AUDPC) calculated to summarise disease progression over time for each individual plant, n=18. Letters represent Tukey HSD statistical significance groupings (p<0.05).

**Supplementary Figure 4:**
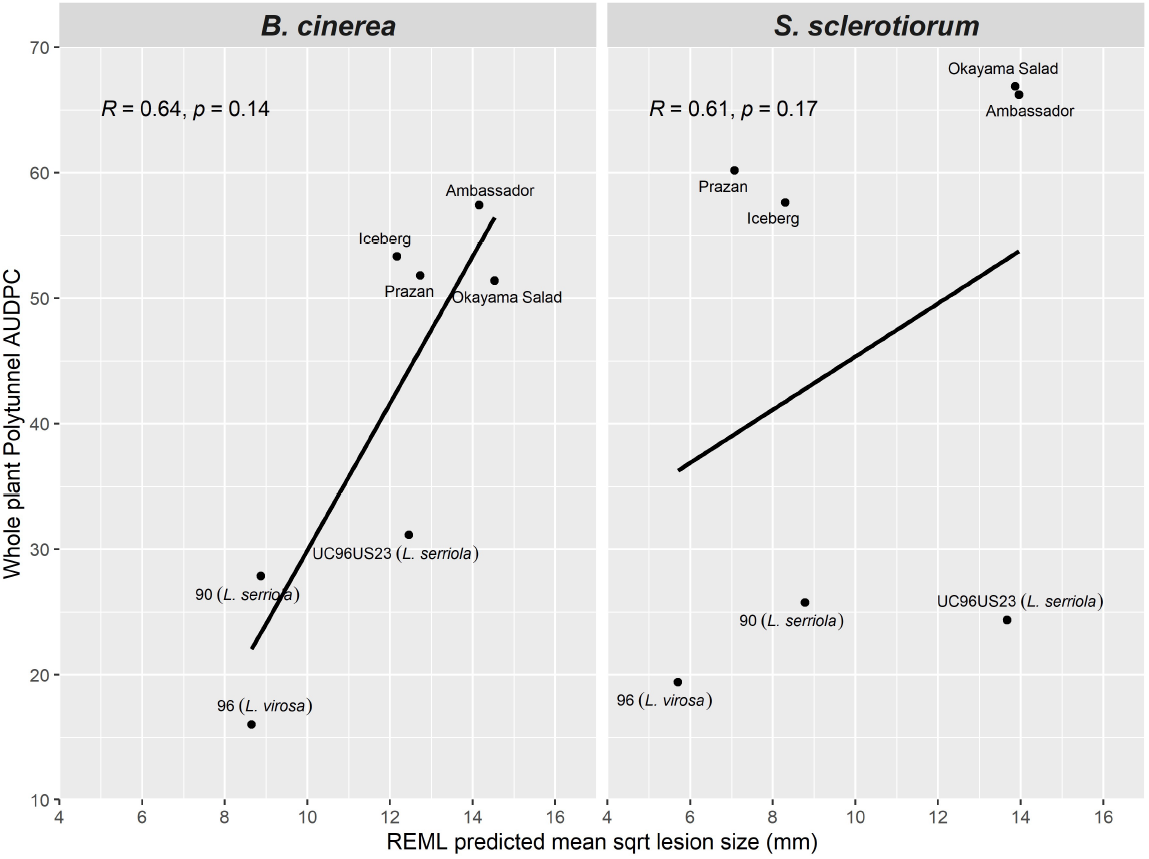
Correlation of REML predicted detached leaf assay square root lesion size (mm) with AUDPC in whole plant inoculations of *B. cinerea* (left) and *S. sclerotiorum* (right) for seven lettuce accessions. Pearson’s correlation coefficient (R) values and p-values are shown. All accessions are *L. sativa* unless otherwise indicated.

**Supplementary Figure 5:**
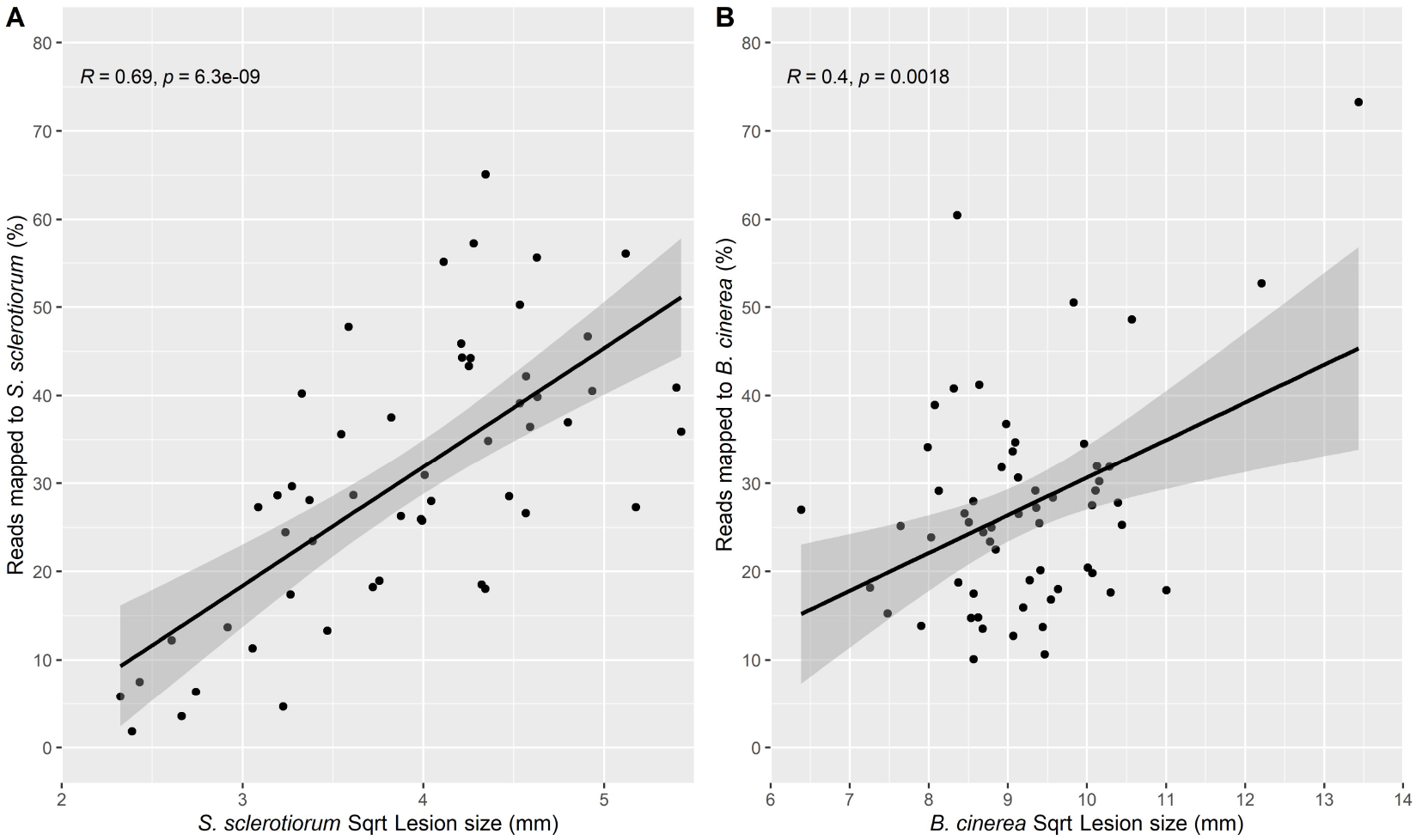
Pearson’s correlation of RNAseq reads in each sample that map to fungal transcripts versus lesion size in (A) *S. sclerotiorum* and (B) *B. cinerea* inoculated samples.

**Supplementary Figure 6:**
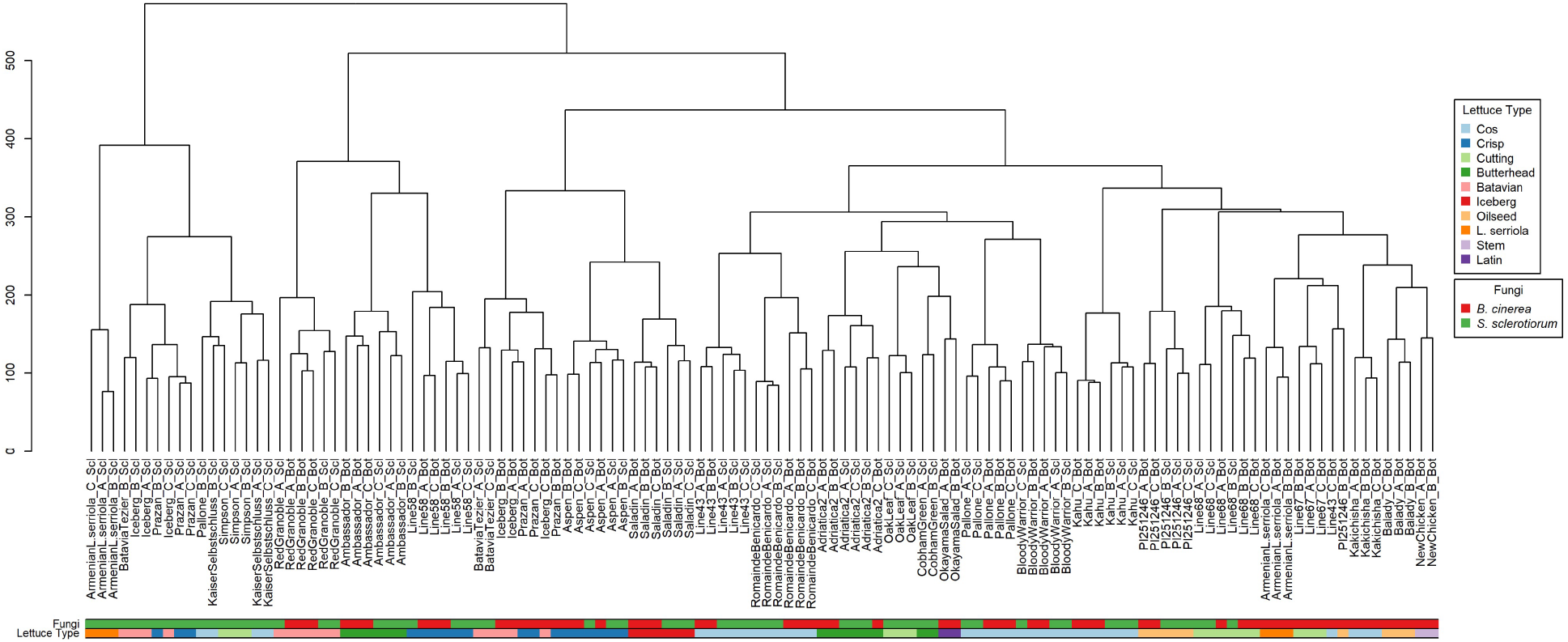
Dendrogram showing Euclidian distance between lettuce diversity set RNAseq samples. With a few exceptions, biological replicates of the same accession cluster together.

**Supplementary Figure 7:**
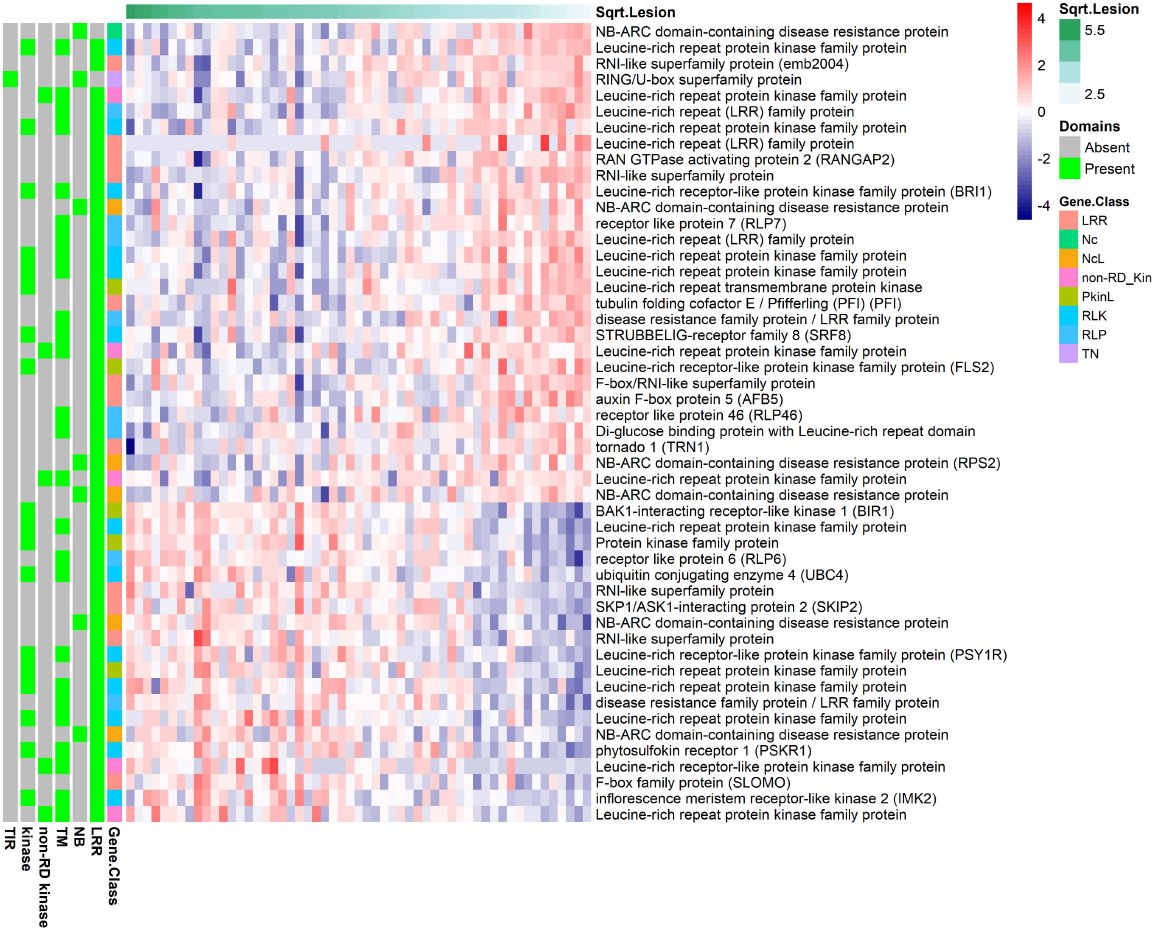
Expression of the top 50 lettuce genes whose expression is correlated with *S. sclerotiorum* lesion size, and which were classified as non-NLR pathogen recognition receptors (Christopoulou et al. 2015 classification indicated in the column gene.class). This classification was based on the presence of any combination of the following domains: leucine-rich repeats (LRR), nucleotide-binding (NB), NB Coiled-coil type (Nc), transmembrane (TM), kinase, non-arginine-aspartate kinase (non-RD kinase) and TOLL/interleukin-1 receptor (TIR). Additional gene nomenclature includes NcL: NC plus L domains; PkinL: kinase plus L; RLK: receptor-like kinase; RLP receptor-like protein. The individual lettuce samples are ordered left to right on the basis of lesion size after inoculation with *S. sclerotiorum*, with the most susceptible (largest lesion size) on the left and most resistant (smallest lesion size) on the right. Log_2_ expression is indicated by the red/blue scale.

**Supplementary Figure 8:**
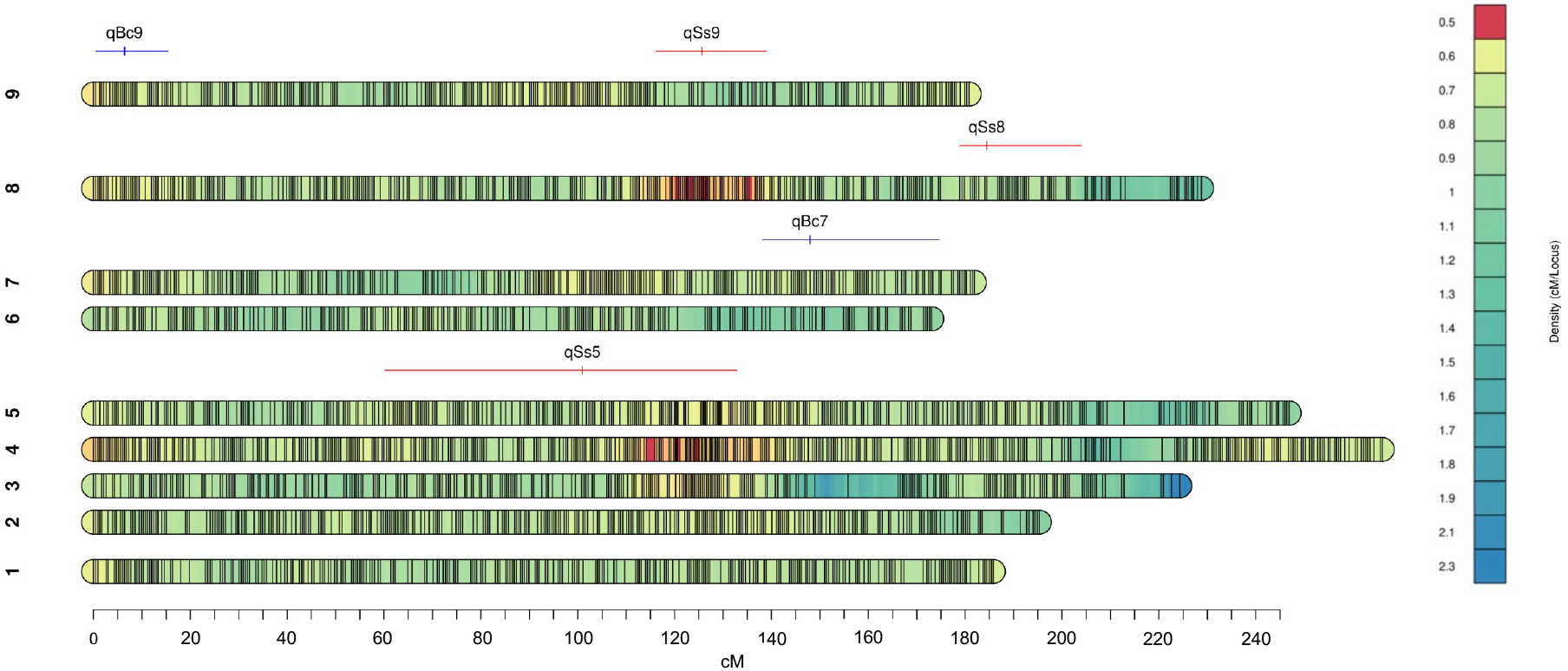
Location of resistance QTL on the Armenian *L. serriola* x PI251246 marker density map. Horizontal bars show the 1.5 LOD confidence interval of the QTL loci and the vertical bar shows the location of the peak QTL marker.

**Supplementary Figure 9:**
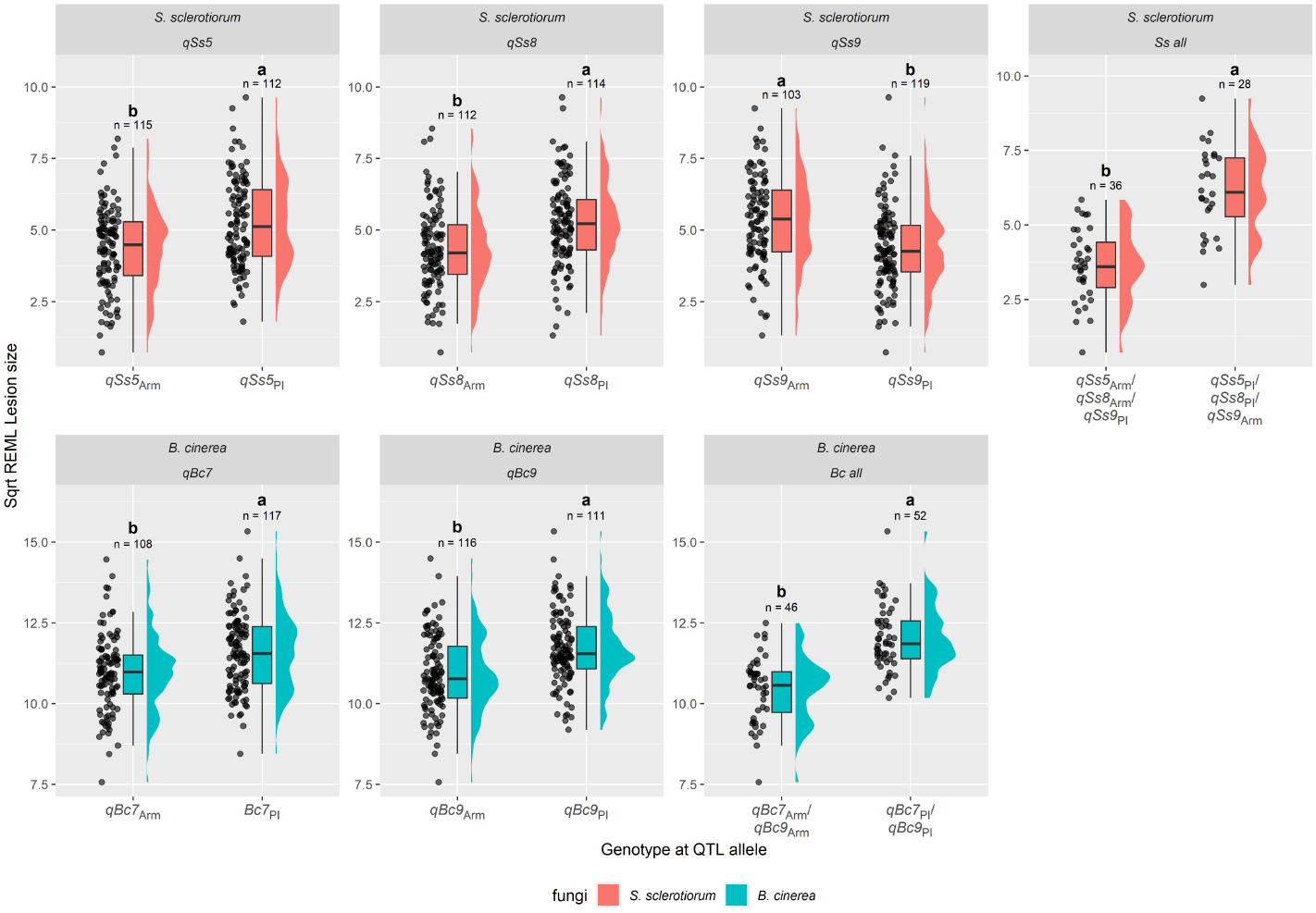
Variation in lesion size between RILs containing alleles originating from PI251246 (PI) or Armenian *L. serriola* (Arm) at identified QTL markers. Violin plots show the distribution of least-squares predicted mean lesion size for RILs after inoculation with the pathogen relevant to the identified QTL (*B. cinerea* blue, *S. sclerotiorum* red). Black horizontal bars represent the mean. Letters show statistical significance groupings (Tukey HSD p<0.05) and the number of samples tested is indicated for each genotype.

**Supplementary Figure 10:**
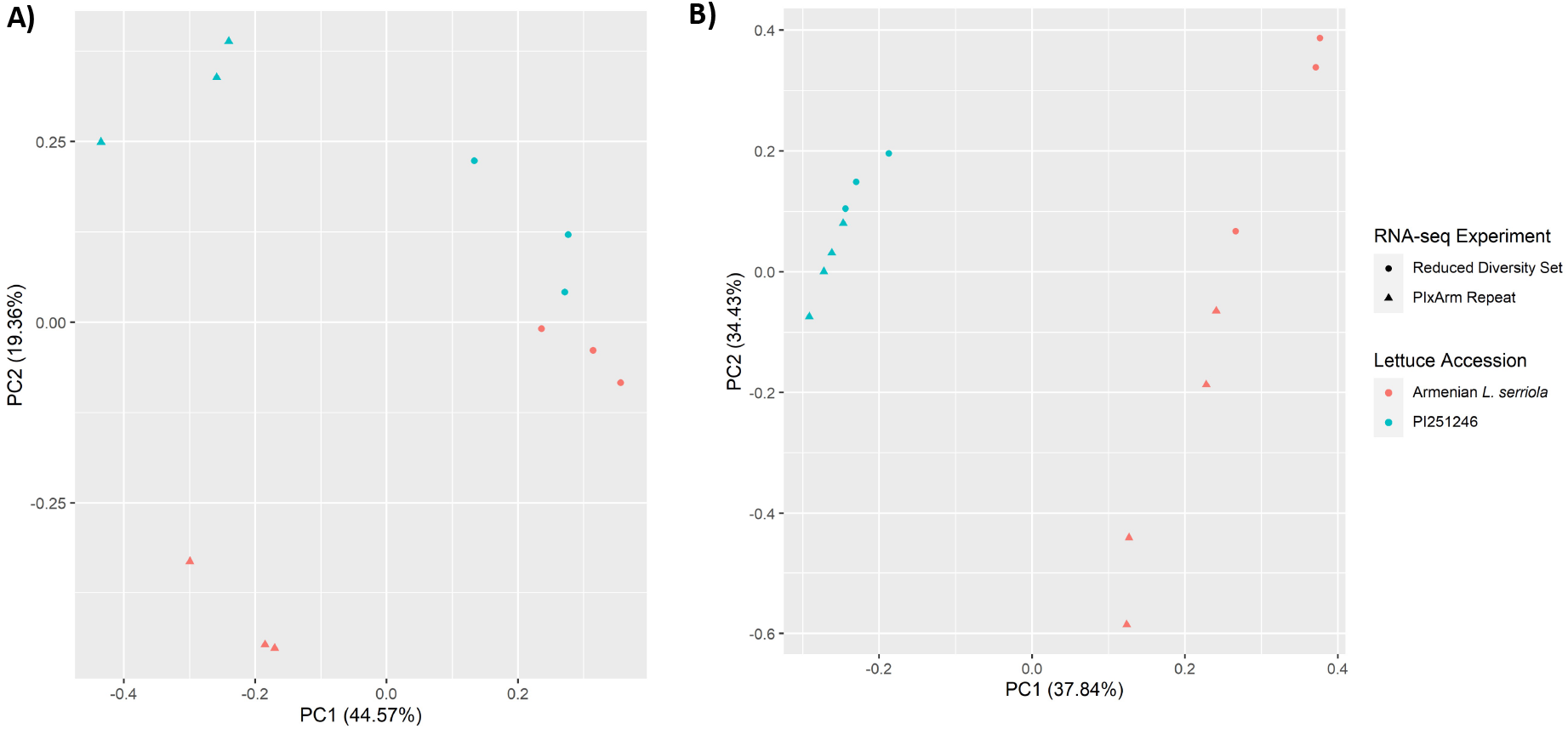
Principal Component Analysis (PCA) of PI251246 (blue) and Armenian *L. serriola* (pink) RNAseq data after pathogen infection with (A) *B. cinerea* and (B) *S. sclerotiorum*. The PCA plot shows two independent experiments: Diversity Set RNAseq (circles) and the mapping population parent repeat (triangles). In *S. sclerotiorum* infected samples, there is a clear separation across PC1 between the parental lines that is consistent across experiments. In *B. cinerea* infected samples, the largest separation across PC1 appears to reflect different experiments, but each parent line clearly separates within the experiment across PC2.

